# 3DeFDR: Identifying cell type-specific looping interactions with empirical false discovery rate guided thresholding

**DOI:** 10.1101/501056

**Authors:** Lindsey Fernandez, Thomas G. Gilgenast, Jennifer E. Phillips-Cremins

**Affiliations:** Department of Bioengineering, University of Pennsylvania, Philadelphia, PA 19104; Epigenetics Institute, Perelman School of Medicine, University of Pennsylvania, Philadelphia, PA 19104; Department of Genetics, Perelman School of Medicine, University of Pennsylvania, Philadelphia, PA 19104

**Keywords:** 3D chromatin looping interactions, higher-order chromatin architecture, epigenetics, Chromosome-Conformation-Capture, chromatin dynamics, false discovery rate

## Abstract

The mammalian genome is connected into tens of thousands of long-range looping interactions critically linked to spatiotemporal gene expression regulation. An important unanswered question is to what extent looping interactions change across developmental models, genetic perturbations, drug treatments, and disease states. Although methods exist for calling loops in single biological conditions, there is a severe shortage of computational tools for rigorous assessment of cell type-specific looping interactions across multiple biological conditions. Here we present 3DeFDR, a simple and effective statistical tool for classifying dynamic looping interactions across biological conditions from Chromosome-Conformation-Capture-Carbon-Copy (5C) data. 3DeFDR parses chromatin contacts into invariant and cell type-specific classes by thresholding on differences in modeled interaction strength signal across two or three cellular states. Thresholds are iteratively adjusted based on a target empirical false discovery rate computed between real and simulated 5C maps. 3DeFDR enables the sensitive detection of high-confidence looping interactions and markedly reduces false positives when benchmarked against a classic analysis of variance (ANOVA) test, our newly formulated parametric likelihood ratio test (3DLRT), and the leading Hi-C differential interaction caller diffHic. 3DeFDR also sensitively and specifically calls loops in Mb-scale genomic regions parsed from Hi-C data. Our work provides a statistical framework and an open-source coding library for identifying dynamic long-range looping interactions in high-resolution 5C data from multiple cellular conditions.

## Introduction

Chromosome-Conformation-Capture (3C)-based molecular techniques have recently been coupled with high-throughput sequencing to generate genome-wide maps of higher-order chromatin folding [1-3]. A number of massively parallel 3C-based technologies query genome folding in a protein-independent manner, including Hi-C, 4C, 5C, and Capture-C [4-10]. All four techniques rely on proximity ligation to convert physically connected chromatin fragments into sequencing-based counts information. Briefly, chromatin is fixed in its native architectural state across a population of cells and then digested with a restriction enzyme. Restriction fragments are ligated to form billions of hybrid ligation junctions between two distal genomic loci. The two fragments in a given ligation junction can then be detected using high-throughput sequencing, and their frequency of ligation is proportional to their spatial proximity across a population of cells. Hi-C detects all chromatin interactions genome-wide using high-throughput sequencing, whereas 5C and Capture-C use tiled probes to selectively sequence large, Mb-scale subsets of the genome. 4C queries all genome-wide contacts for a single, specific restriction fragment. Thus, the protein-independent 3C technologies of Hi-C, 5C, and Capture-C can be used to create high-resolution spatial maps of genome folding on the scale of Mb to genome-wide coverage.

Recently published 3C-based sequencing studies have revealed that the mammalian genome is folded into a hierarchy of distinct architectural features, including: A/B compartments, lamina associated domains (LADs), topologically associating domains (TADs), subTADs, and long-range looping interactions [6, 8, 10-19]. The highest resolution maps to date have enabled the detection of tens of thousands of long-range looping interactions genome-wide [11, 20]. Loops are uniquely different from other forms of long-range chromatin interaction features and are represented in Hi-C maps as groups of adjacent pixels which form a punctate focal increase in interaction frequency enriched above local TAD and subTAD structure [11]. Loops connected by the architectural protein CTCF are thought to create TADs/subTADs that demarcate the search space of enhancers for their target promoters [21-24]. Enhancers loop to promoters via cell type-specific architectural proteins and cohesin to govern spatiotemporally regulated transcription [25-28]. Initial studies have suggested that specific subsets of loops can reconfigure in development, disease, and in response to genetic perturbations [20, 23, 25, 26, 29-35]. Generally, however, it remains unknown to what extent looping interactions are dynamically altered genome-wide as cells switch fate, due in part to the relative paucity of computational methods to evaluate statistically significant changes in looping strength across multiple biological conditions.

As high-resolution Hi-C and 5C chromatin folding maps begin to accumulate in developmentally relevant cellular models, there is an increasing need for methods to (1) sensitively detect long-range looping interactions and clearly distinguish them from other classes of architectural features such as local TAD/subTAD structure and compartments and (2) rigorously classify loops by their dynamic behavior across cell types. To our knowledge, computational tools are not yet available to statistically test looping interactions for their differential signal across two or three conditions in 5C data. A number of computational methods identify loops in individual libraries generated by Hi-C. Bicciato and colleagues performed a detailed comparison of Hi-C loop calling pipelines, including HiCCUPS [36], GOTHiC [37], HOMER (http://homer.ucsd.edu/homer/interactions/), diffHi-C [38], HIPPIE [39], and Fit-Hi-C [40]. The conclusion from this study was that loop calling methods in individual samples exhibit vastly different performance, with no clear gold-standard emerging [41]. Importantly, most loop calling pipelines were developed on low-resolution maps (40 kb up to 1 Mb bins) generated with the first-generation dilution Hi-C experimental procedure. More recently, Hi-C maps have achieved 1-5 kb resolution through higher read depth and markedly improved spatial noise due to second generation in situ ligation and digestion techniques [11, 20]. The emerging model from second-generation Hi-C studies is that quantitative loop detection in individual libraries requires rigorous modeling of local chromatin domain structure. HICCUPS explicitly models and accounts for locus-specific TAD/subTADs, and has therefore emerged as the leading candidate for identifying bona fide looping interactions in individual Hi-C maps.

It has become clear that loops must be distinguished from other architectural features before evaluating their differential signal across cell types. Three tools (diffHic [38], FIND [42], and HiBrowse [43]) have been published to identify generally differential interactions between conditions in Hi-C data. All three in their published, first-generation form were not designed or verified to distinguish loops from higher-order folding patterns such as A/B compartments, TADs, or subTADs [42]. In the absence of accounting for these features, a large proportion of the differential interactions will be due to cell type-specific fluctuations due to technical biases, local chromatin domains, or higher-order compartments. Noteworthy, the diffHic manuscript indicates that modeling local chromatin domain structure would be essential to evaluate cell type-specific loops, suggesting that second-generation tools might be available in the future [38]. Computational tools have also been published to call within- and across-condition loops from libraries generated by Hi-ChIP and ChiA-PET assays [44-49]. However, statistical frameworks built for protein-dependent 3C-methods cannot address the technical challenges unique to 5C and Hi-C data. Overall, a gold-standard statistical methodology for cell type differential loop detection in 5C data is an important unmet need. Moreover, differential loop calling remains an open question for all high-resolution, protein-independent proximity ligation data, and there is no gold-standard differential Hi-C loop calling pipeline published which could be then be easily re-configured to find dynamic loops across multiple conditions in 5C data.

Here, we present 3DeFDR, a new statistical method and coding implementation for identifying cell type-specific looping interactions from 5C data across two or three biological conditions. To facilitate comparison of looping signal across different libraries, 3DeFDR begins by ensuring that all possible genomic interactions have been bias-corrected, modeled with respect to the expected contact frequency due to distance-dependent interactions and local TAD/subTAD structure, assigned p-values, and converted to an interaction score. 3DeFDR uses a thresholding scheme on the change in interaction score signal to stratify seven classes of cell type-specific looping interactions among three biological conditions. By applying the same thresholds to 5C pseudoreplicates simulated from the same biological condition, 3DeFDR computes an empirical false discovery rate (eFDR) under the assumption that all pseudoreplicate contacts passing the thresholds are false positives. 3DeFDR generates pseudoreplicates by modeling the raw 5C count mean-variance relationship and simulating counts for each interaction by parameterizing a negative binomial distribution. To determine the set point of the interaction score thresholds, we provide a controlling procedure in which we iterate thresholds to achieve an *a priori* determined eFDR. We benchmark the performance of 3DeFDR against (i) an established ANOVA test, (ii) our own newly formulated parametric likelihood ratio test (3DLRT), and (iii) the leading published Hi-C non-specific differential interaction calling method diffHic. Cell type-specific loops called by 3DeFDR have fewer false positives and are more strongly enriched for chromatin modifications characteristic of the cellular state in which the loops are present compared to the benchmarking tests. 3DeFDR also has utility in detecting cell type-specific loops in Hi-C data, whereas existing published tools conflate differential signal from many architectural features. 3DeFDR and the parametric benchmarking test 3DLRT are freely availble as Python coding packages to support the next wave of discoveries in dynamic looping.

## Results

We set out to address two critical challenges in the analysis of looping interactions in 5C data. First, there is a paucity of methods for robustly classifying dynamic looping interactions among multiple cellular conditions, a problem which becomes more challenging as the number of conditions increases. Second, there is not yet a gold standard method for determining 5C thresholds to sensitively parse bona fide looping interactions from background non-looping interactions while minimizing false positives. Our goal was to develop a statistical framework and coding implementation to rigorously identify differential looping interactions from 5C maps across two or three conditions using a target FDR to choose thresholds (Figure 1A).

**Figure 1.**
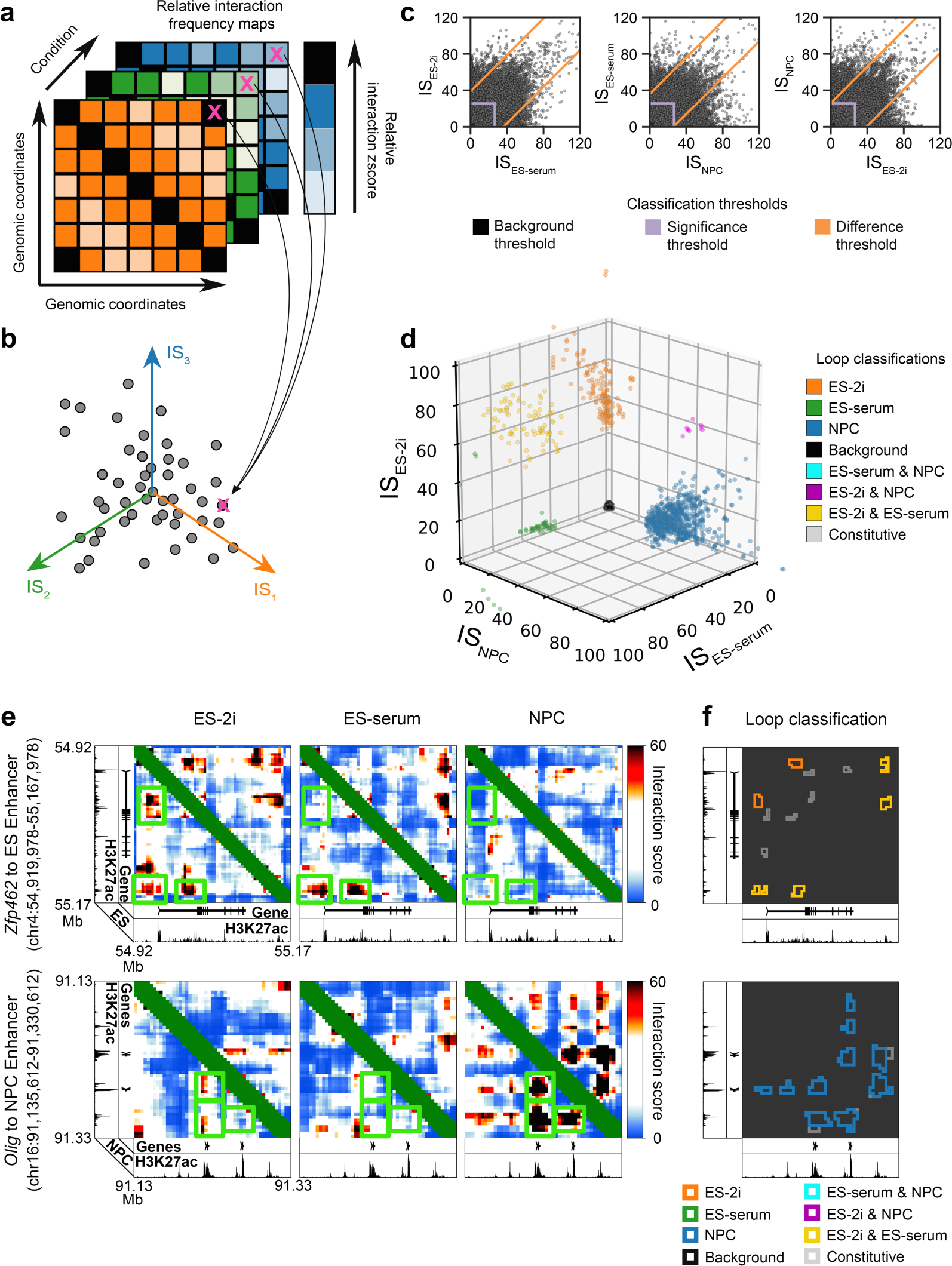
Overview of interaction score thresholding procedure for cell type-specific looping interaction classification. **(A)** A 5C dataset is input as a set of interaction frequency matrices, each matrix capturing the same set of genomic contacts under a different cellular condition. **(B)** Raw 5C counts are converted to interaction scores (IS) which reflect bias-corrected, sequencing depth normalized, local expected background signal normalized, and modeled interaction frequency values that are comparable within and between conditions (detailed in Supplementary Methods and Figure 4). **(C)** ISs are thresholded to allow detection and classification of looping interactions that are significantly differential across cellular conditions. **(D)** Seven looping interaction classes after a 3-way thresholding scheme on ES-2i, ES-serum, and NPC cellular states. **(E)** IS heatmaps at two selected genomic loci. Green boxes highlight regions of qualitatively apparent differences in looping signal. **(F)** Loop classification results after applying 3DeFDR’s 3-way IS thresholding procedure.

Here we develop, apply, and benchmark 3DeFDR using 5C data across three distinct cellular states: embryonic stem (ES) cells cultured in 2i/LIF media representing the naive pluripotent state, ES cells cultured in LIF/serum representing the primed pluripotency state, and primary neural progenitors isolated from neonatal mice representing a multipotent adult stem cell state in the neuroectoderm lineage (**Supplementary Table 1**) [25]. These particular 5C datasets represent large-scale, 4 kb-resolution maps capturing 8 Mb of genomic sequence around key developmentally regulated genes. 5C relies on a primer-based hybrid capture step to selectively detect ligation junctions across specific genomic regions, thus enabling the creation of high-resolution matrices with a strikingly lower number of reads (∼30-40 Million per sample) compared to Hi-C (∼3-6 Billion per sample). We have recently determined that looping interactions are markedly reconfigured during the transition from naive pluripotency to multipotency, thus making this data ideal for the testing and development of our statistical framework. We test and validate 3DeFDR with a three cellular state experiment, but the statistical framework and code is set up to analyze a two cellular state experiment as well.

We first started by modeling and correcting biases, artifacts, and local chromatin domains in individual replicates. Despite their nuanced technical differences, data from protein-independent proximity ligation techniques share several common features, including: (1) distance-dependent background interaction signal in which non-specific interaction frequency is highest for the closest fragment-fragment pairs on the linear genome and decays as the distance separating the genomic fragments increases [6], (2) biases in ligation and amplification frequency caused by GC content and length of restriction fragments [50, 51], (3) library complexity and sequencing depth differences across independent experiments for the same biological sample leading to nonlinear batch effects [52], and (4) highly locus-specific structure due to higher-order folding of chromatin into TADs, subTADs, and compartments [11]. One must model and address these features to ensure a rigorous analysis of looping interactions.

We reasoned that a differential loop calling method would have the most utility across protein-independent proximity ligation data if it started with a modified interaction score (IS) in which all confounders and background signal had been corrected. We recently discovered that sequence-related biases are not constant across cell types and replicates. Therefore, as is routinely done with Hi-C data, we used matrix balancing to correct for fragment-specific biases caused by GC content and restriction fragment length for every replicate individually (**detailed in Supplementary Methods**). We also used conditional quantile normalization to normalize all replicates for non-linear library complexity and sequencing depth differences (**detailed in Supplementary Methods**). It is also widely known that the distance-dependent background signal and local chromatin domain structure are widely variable across cell types and highly unique to each genomic region. Thus, we used the donut and lower left filters [11, 53] to model the distance-dependent and TAD/subTAD expected background signal for every interaction in the genome and every replicate individually (detailed in **Supplementary Methods**). After bias correction, background normalization, and expected modeling, we assign pvalues to every pixel in the 5C heatmap and compute an interaction score (IS) that allows for direct comparison of each bin-bin pair across replicates and conditions (Figure 1B). Moreover, the use of modeled IS as the random variable for differential testing allows 3DeFDR to have utility for matrices of any protein-independent 3C-based data that has been bias corrected, normalized, modeled, and transformed into p-values using analysis techniques taylored to the specific method.

To identify differential looping interactions, we use a classification technique that relies on three-way thresholding on the difference in IS across cellular conditions (Figure 1C**, Supplementary Figure 1, Supplementary Table 2**). For each biological replicate, we begin with a framework in which matrix *IS* is a square, symmetric array of interaction scores from a modeled and bias-corrected 5C experiment. The matrix *IS* has dimensions n by n, where n is the number of genomic bins at any particular genomic region, *r*. Thus, 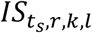 represents the interaction score between genomic bins *k* and *l* in region *r* as recorded for biological replicate *s* of condition *t* (detailed in **Supplementary Methods**). We first identify potential looping interactions by parsing only bin-bin interactions with an 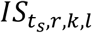 greater than a specific significance threshold *g* for all replicates in at least one condition (purple lines, Figure 1C). We then apply a series of thresholds (orange lines, Figure 1C) on the difference in 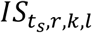 across all three cellular conditions (**Supplementary Figure 1E-G, Supplementary Table 2**). To ensure the most conservative estimate of looping classes, we apply the thresholds on the minimum difference in IS across replicates of each condition. Thus, the end result is a preliminary set of seven classes of looping interactions: (1) ES-2i only, (2) ES-serum only, (3) NPC only, (4) ES-2i & ES-serum only, (5) ES-2i & NPC only, (6) ES-serum & NPC only, (7) constitutive across all three cell types (Figure 1D). Examples of ES-2i only, ES-2i & ES-serum only, and NPC only interactions are illustrated in Figure 1E-F.

We next used estimation and control of an empirical false discovery rate (eFDR) to guide the final placement of the difference thresholds for each looping class (orange lines, Figure 1C, detailed in **Supplementary Methods)**. The false discovery rate (FDR) is by definition *FDR* = *E*(*V*/*R*) where *V* is the number of false positives among tests declared significant and *R* is the total number of tests declared significant. Here, *R* is trivial to compute from our set of three conditions (*T* = {*A*, *B*, *C*} where A is ES-2i, B is ES-serum, and C is NPC) and six replicates (*S* = {*A*1, *A2*, *B*1, *B2*, *C*1, *C2*}) as the total number of significant bin-bin interactions in a given looping class (*H* = {{*A*}, {*B*}, {*C*}, {*A*, *B*}, {*A*, *C*}, {*B*, *C*}}). However, *V* is not known and requires a method for estimating the false positive rate of our three-way thresholding procedure.

We hypothesized that *V* is approximately equal to the total number of interactions incorrectly labeled differential when applying 3DeFDR to the set of biological samples with no true differential loops. We defined our null dataset as a set of samples that are replicates of a single cellular condition but assigned a random set of labels matching conditions *T*. Our key assumption in formulating this approach is that that the false positive rate (FPR) of calls on the null dataset (*FPR* _*null*_) is approximately equivalent to that of the experimental dataset (*FPR* _*exp*_), such that *FPR* _*null*_ ≈ *FPR* _*exp*_. We computed and controlled an empirical false discovery rate (eFDR) as in Equation 1:

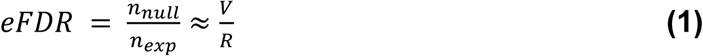

where *n* _*exp*_ is the total number of interactions classified as significantly differential for a particular looping class using the experimental conditions *T* and *n* _*null*_ is the total number of interactions classified as significantly differential in the null dataset, which approximates *FPR* _*exp*_.

It is often cost prohibitive to generate six biological replicates of 5C data for each condition. Therefore, we generated 5C replicate simulations to populate the null sample set. We simulated 5C replicates of the same condition at the level of fragment-level counts after conditional quantile normalization (i.e. 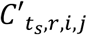 interactions for every *i* ^*th*^ and *j* ^th^ fragment for each genomic region *r* after quantile normalization and before binning and matrix balancing). Our rationale for this decision is that library complexity, batch effect and sequencing depth terms would not have to be explicitly included in our count generating models. To construct our simulation generating model, we first computed the sample mean and sample variance for every interaction in every condition (Equations 2-3):

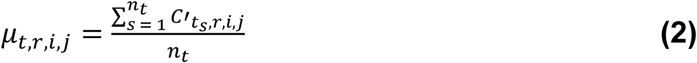

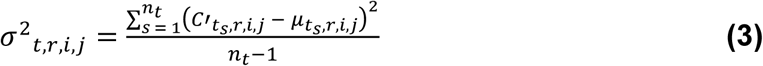

where *n* _*t*_ is the number of replicates of condition *t* and 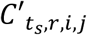 is the conditional quantile normalized raw 5C counts of interaction (*t*, *r*, *i*, *j*) in the *s* ^*th*^ replicate of condition *t* for every *i* ^*th*^ and *j* ^*th*^ fragment ligation in genomic region *r*.

Most genomics experiments suffer from poor parameter estimation due to the low number of replicates that are financially and logistically feasible to generate for every biological condition. To improve parameter estimates, we modeled the mean-variance relationship (MVR) between *μ* _*t,r,i,j*_ and *σ* ^2^ _*t,r,i,j*_ by pooling all interactions in all regions (Figure 2). We stratified quantile normalized counts, 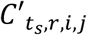, for all regions by their linear genomic distance using a dynamic size window (Figure 2A). For distance regime 1 (0-150 kb), we stratified the interactions into fine-grained, 12 kb-sized sliding windows, w _0-150kb_, with a 4 kb step. For distance regime 2 (151-600 kb), we stratified the interactions into 24 kb-sized sliding windows, w _151-600kb_, with an 8 kb step. For distance regime 3 (601-1000 kb), we stratified the interactions into coarse-grained, 60 kb-sized sliding windows, w _601-1000kb_, with a 24 kb step. We found that the variance was greater than the mean across all genomic distance scales, indicating that 5C counts data are overdispersed (Figure 2B). For each window in each distance regime, we modeled the MVR by fitting the function (Equation 4):

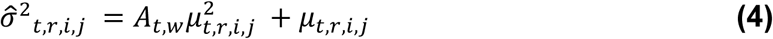

to all *μ* _*t,r,i,j*_ and *σ* ^2^ _*t,r,i,j*_ to find the overdispersion parameter, *A* _*t,w*_, at each distance scale (detailed in **Supplementary Methods**). We found that *A* _*t,w*_ also varied as a function of distance and was unique to each cell type (Figure 2C-D). Together, these data demonstrate that 5C counts are overdispersed and that the overdispersion parameter varies as a function of distance and cellular state.

**Figure 2.**
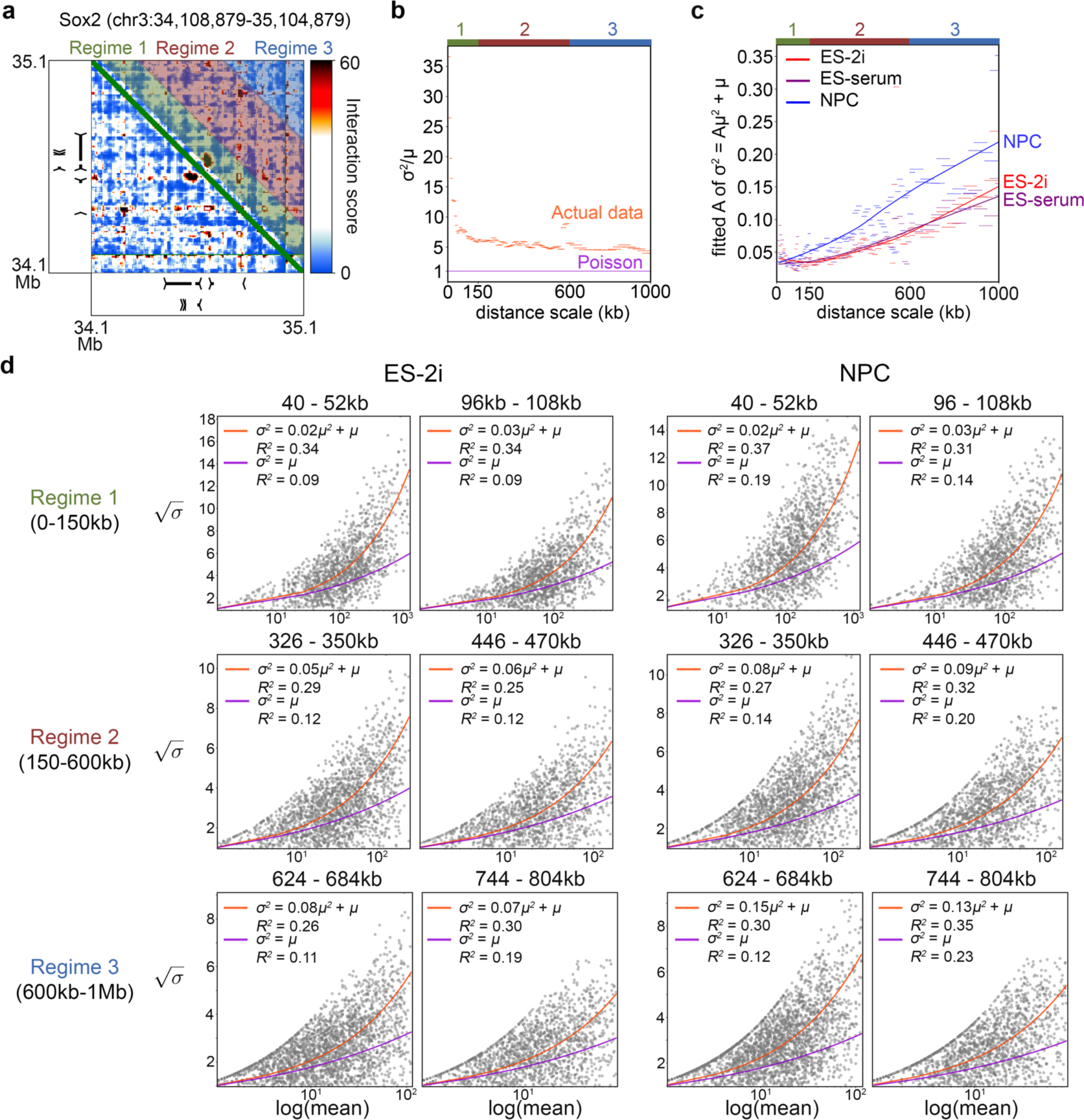
5C counts are overdispersed and their mean-variance relationship varies as a function of linear genomic distance and cellular condition. **(A)** Raw 5C contacts are stratified by genomic distance prior to characterization of their mean-variance relationship. In each of three regimes, the width of these windows is determined using a different binning scheme. **(B)** The coefficient of variation for raw 5C counts are plotted against genomic distance for each sliding window’s median interaction distance. Each window captures counts from all genomic regions in the dataset in the ES-2i condition. **(C)** The dispersion parameter, A, for each distance scale window is computed by fitting sample means and variances to the function σ ^2^ = A*µ ^2^ + µ. Dispersion vs. distance scale trend lines were generated by Loess smoothing. **(D)** Mean-variance models are displayed for representative genomic distance windows from all three distance regimes. Fits of the Poisson mean-variance relationship (σ ^2^ = µ) and the negative binomial mean-variance relationship (σ ^2^ = A*µ ^2^ + µ) are shown with R ^2^ goodness of fit values.

To generate simulated 5C libraries, we weighted the predicted variance 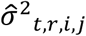 against the original observed variance *σ* ^2^ _*t,r,i,j*_ to generate a final weighted variance 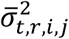 for each interaction at each distance scale as in Equation 5 (detailed in the **Supplementary Methods)**:

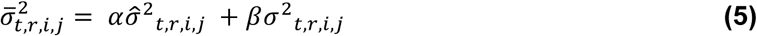

We used *α* = *β* = 0.5 to achieve simulated 5C counts with pairwise correlations on par with that of real replicates while improving the quality of our variance estimate with the predicted contribution (**Supplementary Table 3**). Finally, we parameterized the negative binomial model for each *C*′ _*t,r,i,j*_ interaction and generated simulated counts from our models for each (*t*, *r*, *i*, *j*) interaction (Equation 6):

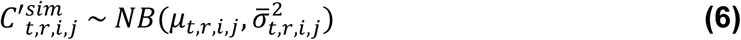

A simulated replicate is created by drawing counts for each (*t*, *r*, *i*, *j*) interaction from a random point on its respective parameterized negative binomial distribution. We then subjected the simulated 5C libraries, 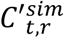, to the same matrix balancing, binning, expected normalization and modeling as the real 5C libraries (**Supplementary Methods**). Simulated 5C counts were highly similar to real 5C data in a qualitative comparison (Figure 3A-D**, Supplementary Figure 2**). Moreover, for the final predicted variance estimates (Equation 5 weighted at *α* = *β* = 0.5), our simulated 5C libraries exhibit Spearman’s correlations within and between conditions that are nearly equivalent to real replicates (Figure 3E). Together, these data show that 5C libraries can be simulated with a negative binomial distribution parameterized with an overdispersed interaction- and region-specific MVR.

**Figure 3.**
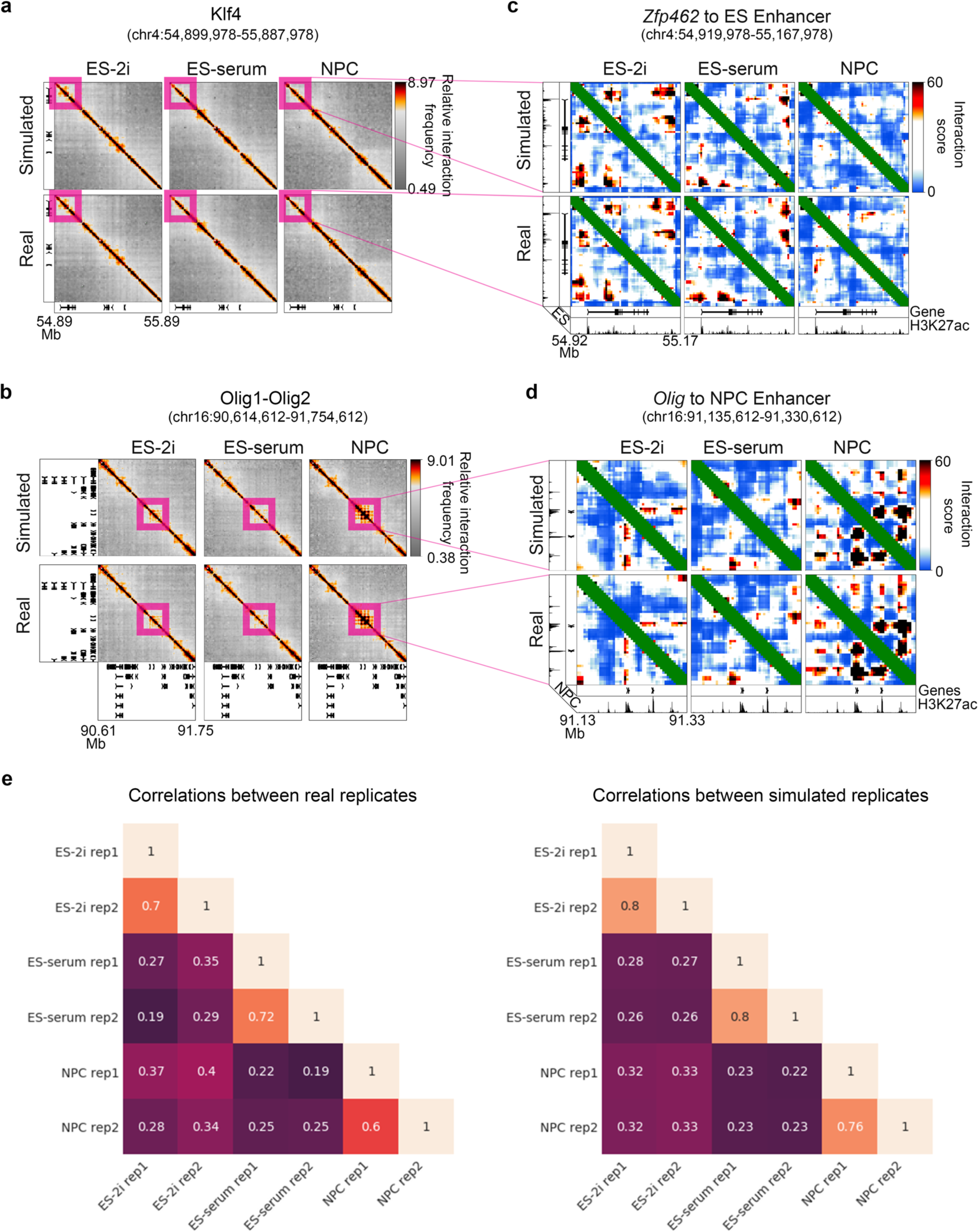
Simulated 5C datasets exhibit strong similarity to experimental 5C datasets. **(A-B)** Heatmaps of relative 5C interaction frequency in the genomic regions surrounding the *Klf4* and *Olig1/2* genes are shown for simulations and real experimental data.**(A)** *Klf4*. **(B)** *Olig1/2*. **(C-D)** Heatmaps of interaction scores in the genomic regions surrounding the *Klf4* and *Olig1/2* genes are shown for simulations and real experimental data. **(C)** *Klf4*. **(D)** *Olig1/2*. **(E)** Matrices of pairwise Spearman’s correlations between real and simulated 5C replicates after conditional quantile normalization (see **Supplementary Methods**).

We next used simulated IS matrices (Figure 4A) to compute an empirical FDR (eFDR) on our looping classes across a sweep of IS difference thresholds applied to both real 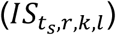 and simulated 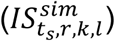 values. For each loop classification, we compute an eFDR for a sweep of difference threshold values, acquiring a difference threshold to eFDR mapping for each class, *eFDR* _*d,h*_, as in Equation 7:

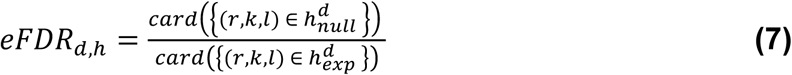

where the numerator is the number of loops assigned to differential class *h* in six replicates of the simulated null dataset at difference threshold *d*, and the denominator is the number of loops assigned to the same class in the experimental dataset at the same difference threshold. We selected our final eFDR threshold *τ* as the location where the eFDR vs. IS difference threshold curves transition from exponentially decreasing to linear (**Supplementary Figure 3**). In this study, *τ* = 1% (Figures 4-5). We performed the eFDR controlling procedure for every differential looping class across our three cellular states (Figure 4B-C). At a 1% eFDR, we identified 1549 constitutive, 152 ES-2i only, 0 ES-Serum only, 299 NPC only, 112 ES-2i & ES-Serum, 0 ES-2i & NPC, and 17 ES-Serum & NPC interacting bin-bin pairs. After clustering bin-bin pairs of a similar looping class by spatial adjacency (**Supplementary Methods**), the end result was 108 constitutive, 15 ES-2i only, 25 NPC only, 10 ES-2i & ES-Serum, and 1 ES-Serum & NPC looping clusters (Figure 4D-E). The 3DeFDR algorithm is designed so that the user can tune the final looping classifications to a pre-determined eFDR.

**Figure 4.**
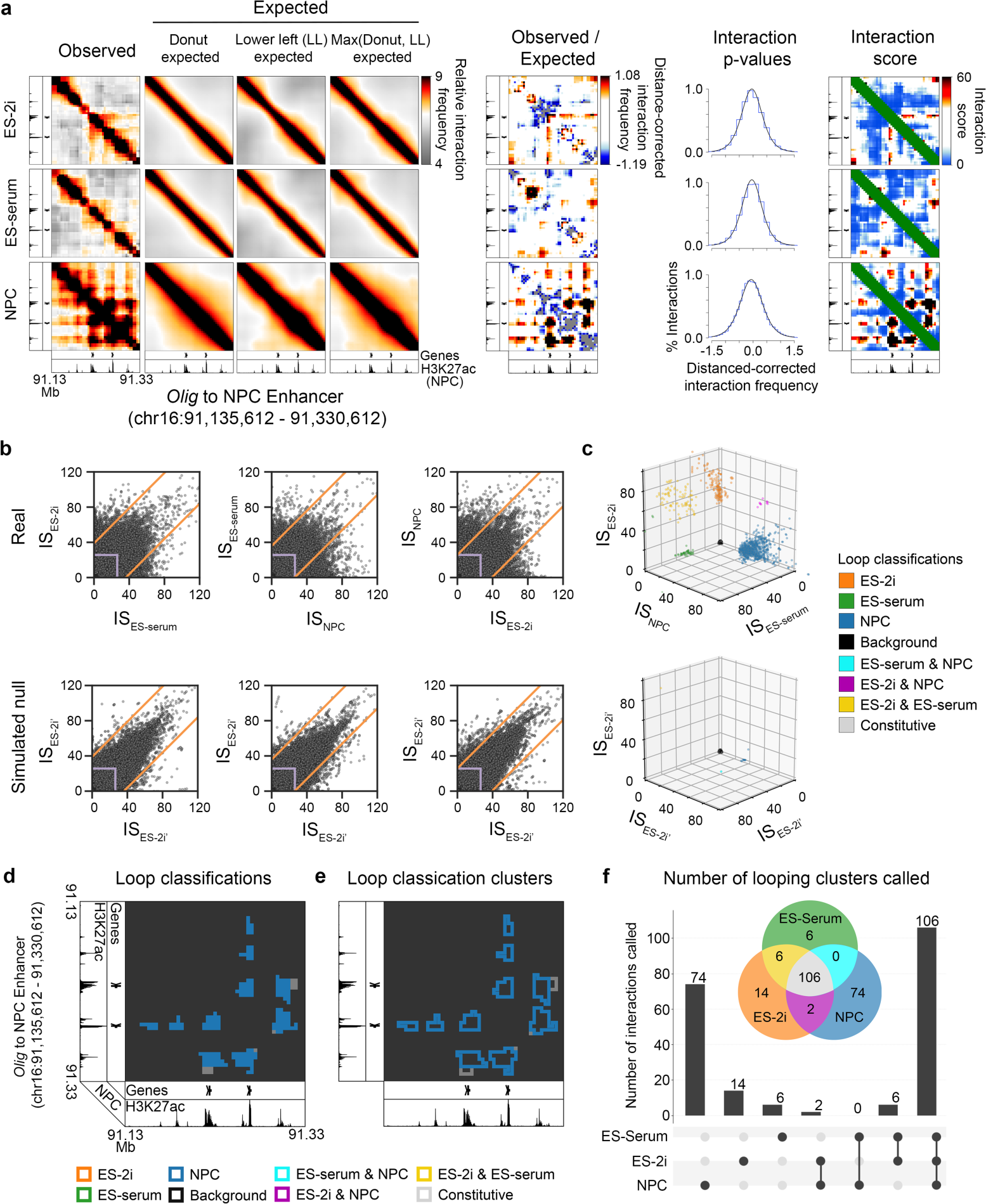
Application of 3DeFDR to find cell type-specific looping interactions across three cellular states. **(A)** Heatmaps representing binned, matrix balanced 5C counts (Observed) around a known looping interaction between the *Olig1* gene and an NPC-specific enhancer (chr16: 91,135,612-91,330,612). Observed counts are divided by the computed local expected signal to obtain background normalized counts (Observed/Expected). These counts are fit to a logistic distribution and the resulting p-values transformed into interaction scores, where interaction score = −10*log2(p-value). **(B)** Interaction scores are thresholded to isolate contacts that are differentially looping across cellular conditions and have signal that meets a baseline requirement for significance. This thresholding procedure is applied to both real and simulated null replicate sets to compute an eFDR. The dynamic thresholding procedure is applied with increasing stringency until a user-specified target false discovery rate is reached. **(C)** Loop classifications obtained with 3DeFDR in real and simulated null replicate sets shown in an interaction scatterplot representation. **(D-E)** Heatmap of final loop classifications at **(D)** individual bin-bin pairs and **(E)** adjacent, similarly classed looping clusters after applying 3DeFDR at a threshold of 1%. **(F)** UpsetR scalable Venn diagrams after 3DeFDR is applied with an eFDR target of 1%.

**Figure 5.**
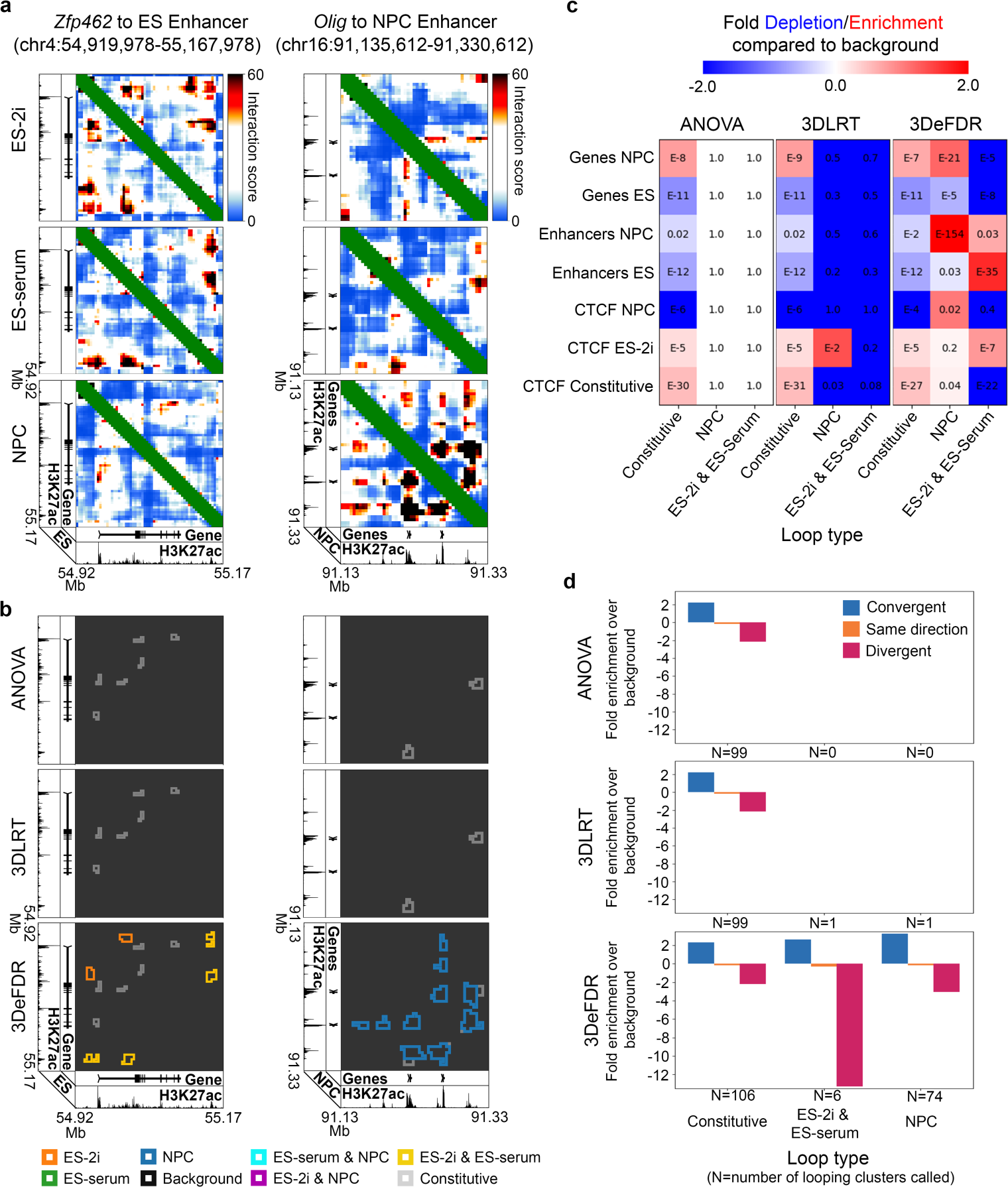
Dynamic 3D chromatin looping interactions identified using 3DeFDR, 3DLRT, and ANOVA. **(A)** Reference interaction score heatmaps for two sample loci. **(B)** Loop classification results achieved with each differential looping detection method at a target false discovery rate (FDR) of 1%. **(C)** Enrichment of cell-type specific markers in loops classified as NPC or ES-2i & ES-serum for each of the three methods at a target FDR of 1%. **(D)** Log fold-change in percent CTCF orientation among loops classified as constitutive, ES-2i & ES-serum, or NPC, over percent CTCF orientation among loops classified as background.

To evaluate the performance of the 3DeFDR pipeline, we implemented two additional methods for classifying differential looping interactions: ANOVA-BH and 3DLRT-BH (**Supplementary Figure 4**). These methods use ANOVA and our newly formulated likelihood ratio test (3DLRT), respectively, to assign a differential looping p-value to every bin-bin pair in an experimental dataset (detailed in **Supplementary Methods)**. In both approaches, output p-values are then corrected for multiple testing using the Benjamini-Hochberg step-up procedure for controlling FDR. When we compared ANOVA and 3DLRT benchmarking tests to 3DeFDR, we found that the three different methods had different optimal FDR thresholds for identifying differential looping structure (**Supplementary Figures 5-7, 9-11**), with 3DeFDR identifying the known, previously reported looping interactions at significantly lower FDR estimates than the other two approaches (Figure 5A-B). Thus, 3DeFDR can identify known cell type-specific looping interactions with a lower false discovery rate than ANOVA and 3DLRT benchmarking tests.

To further understand the dynamic loops called by 3DeFDR, we also compared them to chromatin modifications on the 1-D genome as well as to the performance of the leading non-specific differential interaction caller built for Hi-C data. We observed that classes of differential loops identified by 3DeFDR at an FDR of 1% were strongly enriched for genes and enhancers characteristic of cell types matching those of the identified loops (Figure 5C, **Supplementary Figures 8, 12**). Morever, we observed that convergently and divergently oriented CTCF motifs were enriched and underenriched, respectively, at the base of loops identified by 3DeFDR (Figure 5D). We also observed that diffHic [38] called cell type-specific interactions throughout 5C regions irrespective of whether or not the interactions were loops at both a matched 1% FDR and matched number of signifcant contacts (**Supplementary Figure 14**). Together, these data indicate that 3DeFDR successfully identifies cell-type specific loops at a significantly lower FDR estimate compared to three benchmarks: (1) ANOVA, (2) our own formulated 3DLRT parametric test, and (3) the non-specific Hi-C differential caller diffHic.

Finally, we wondered if 3DeFDR might have utility in calling loops in Hi-C data. Because many biologists are specifically interested in studying a specific genomic locus, we reasoned that 3DeFDR might be effective after stratifying a large Mb-scale matrix from the genome-wide Hi-C matrix. We re-analyzed published ultra-high-resolution Hi-C data from mouse ES cells and ES-derived NPCs from the Cavalli lab [54] (**Supplementary Methods**). We stratified a 4 Mb matrix around *Sox2* and ran the full 3DeFDR pipeline at this locus in the two cell types (Figure 6). Three key loops previously reported in the literature are highlighted on both Observed and Interaction Score matrices, including the ES-specific loop connecting the *Sox2* gene to its ES-specific enhancer (box 1), and the longer-range ES-specific, NPC-specific, and constitutive loops around *Sox2* at this locus (box2, box3) (Figure 6A-B). Similar to 5C analyses, we found that Hi-C data were overdispersed (Figure 6C-D). 3DeFDR identified differential loops at this locus and appropriately classified the invariant, ES-specific, and NPC-specific interactions at an eFDR threshold of 10% (Figure 6E-F). We also applied diffHic [38] to the same Hi-C data. Similar to our observations with 5C (**Supplementary Figure 14**), we observed that diffHic called cell type-specific interactions throughout Hi-C data irrespective of whether or not the interactions were loops at both matched FDR and matched number of signifcant contacts (**Supplementary Figure 13**). Together, these data suggest that 3DeFDR will be useful in the future for calling dynamic loops from stratified regions out of larger Hi-C data sets.

**Figure 6.**
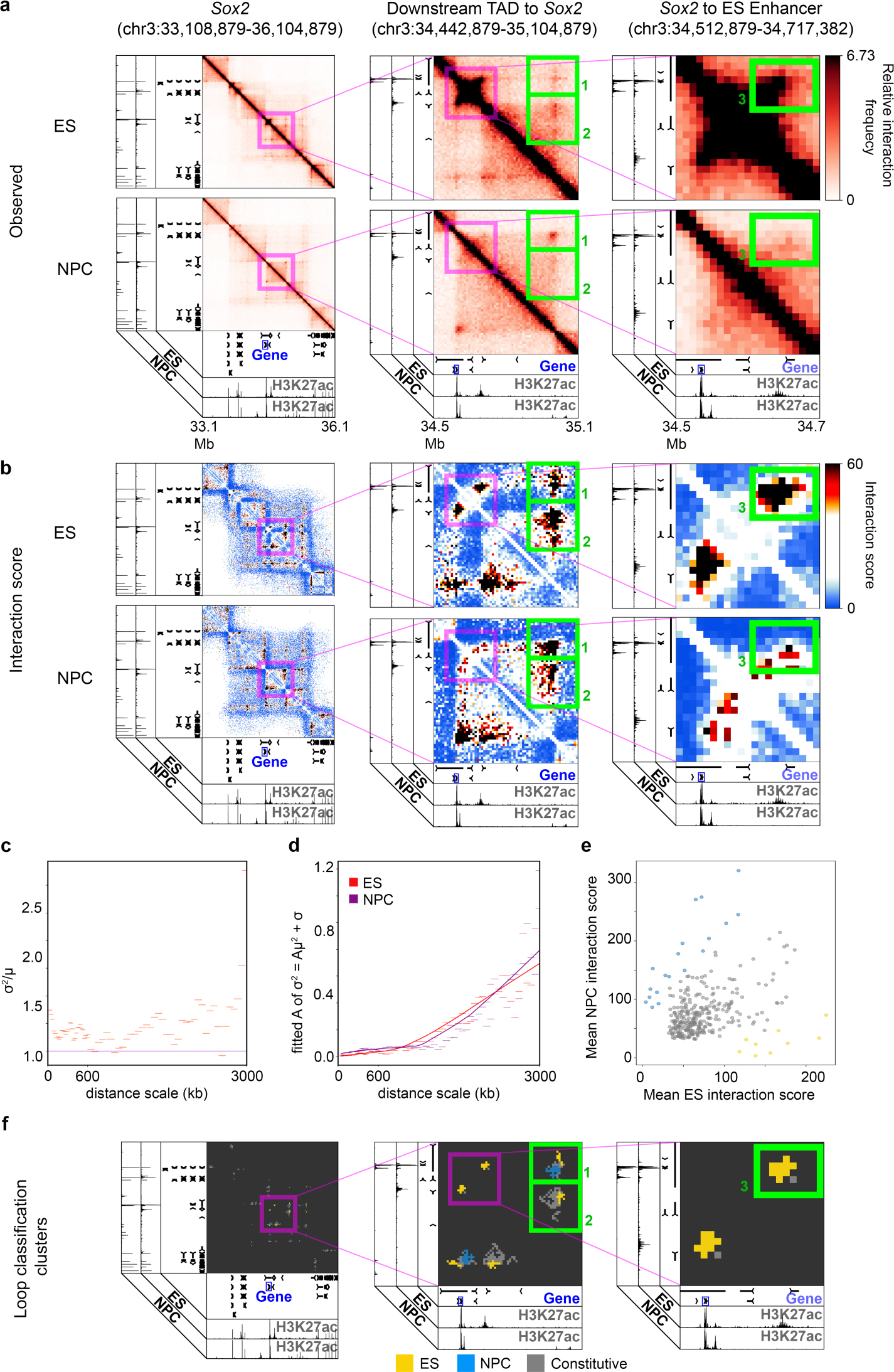
Cell-type specific looping interactions identified from Hi-C using 3DeFDR. **(A)** Reference heatmaps of relative interaction frequency (Observed) Hi-C for *Sox2* and two zoom-in views of the downstream TAD to *Sox2* and its ES enhancer. Boxes 1, 2, and 3 highlight areas of differential looping. **(B)** Reference interaction score heatmaps of the same genomic regions shown in (A). **(C)** The coefficient of variation for observed Hi-C counts are plotted against genomic distance for each sliding window’s median interaction distance. Each window captures counts from all genomic regions in the dataset in the ES condition. **(D)** The dispersion parameter, A, for each distance scale window is computed by fitting sample means and variances to the function σ ^2^ = A*µ ^2^ + µ. Dispersion vs. distance scale trend lines were generated by Loess smoothing. **(E)** Loop classifications obtained with 3DeFDR in real replicate sets shown in an interaction scatterplot representation. **(F)** Heatmaps of final loop classifications for each genomic region at adjacent, similarly classed looping clusters after applying 3DeFDR at a target eFDR threshold of 10%.

## Conclusion

Since the invention of 5C and Hi-C technologies, the field has been in need of statistical methods and computaional tools for identifying differential long-range looping interactions among biological conditions. To date, there is a severe lack of differential loop calling methods available for analysis of 5C data by the scientific community. Moreover, although a small number of ‘general differential interaction identification’ methods have been published for Hi-C data, differential loop calling largely remains an open question because (1) currently available tools do not account for local distance-dependent background signal and TAD/subTAD/compartment structure to identify changes specifically at loops, and (2) Hi-C data at the resolution necessary for looping interaction analysis have only very recently become available.

Identifying bona fide looping interactions from proximity ligation-based data remains a complex problem with no gold standard solution. Moreover, the detection of dynamic looping interactions that exhibit differential signal across biological conditions remains an unsolved technical challenge. Here we provide 3DeFDR as a new statistical framework and computational tool for detecting and classifying differential looping interactions in high-resolution, multi-condition 5C datasets. We note that the performance of 3DeFDR is highly dependent on the quality of the input dataset and how effectively the raw sequencing counts of detected interactions have been processed to reduce batch effects, correct for bias, and account for distance-dependent and TAD/subTAD background signal. We provide 3DeFDR as a modular coding package that the user may integrate into their own 5C or Hi-C analysis pipeline. For the convenience of users, this package includes companion visualization tools for assessing 3DeFDR results to determine how effectively counts have been modeled for simulation, viewing differential loop calls as color-coded clusters, and computing the enrichment of classical epigenetic marks within classes of called loops.

## Methods

Detailed Methods are provided in the **Supplementary Methods**.

## Data availability

Code, sample input data and usage instructions are provided to the Reviewers in a zipped directory and will be freely available via bitbucket upon publication. The data analyzed in this study are summarized in **Supplementary Tables 1, 4, and 5**.

## Acknowledgements

JEPC is a New York Stem Cell Foundation (NYSCF) Robertson Investigator and an Alfred P. Sloan Foundation Fellow. This work was funded by The New York Stem Cell Foundation (J.E.P.C), the Alfred P. Sloan Foundation (J.E.P.C), the NIH Director’s New Innovator Award (1DP2MH11024701; J.E.P.C), a 4D Nucleome Common Fund grant (1U01HL12999801; J.E.P.C) and a joint NSF-NIGMS grant to support research at the interface of the biological and mathematical sciences (1562665; J.E.P.C). This material is based upon work supported by the National Science Foundation Graduate Research Fellowship under DGE-1321851 (L.R.F.) and by an NIH training grant to the University of Pennsylvania (5T32HL007954-18; T.G.G.).

