## Supplementary Materials for "3DeFDR: Identifying cell type-specific looping interactions with empirical false discovery rate guided thresholding"

### SUPPLEMENTARY METHODS

#### 5C data

5C libraries generated with a single alternating primer design[1] in embryonic stem (ES) cells cultured in 2i media (ES-2i), ES cells cultured in serum/LIF (ES-Serum) and primary neural progenitor cells (NPCs) were downloaded from GEO (**Supplementary Table 1**).

#### Hi-C data

Hi-C libraries were downloaded from GEO (**Supplementary Table 5**).

#### 5C data processing pipeline

##### Overview

Raw 5C counts were subject to our previously published 5C count modeling methods[1, 2]. The processing steps described briefly below ultimately resulted in the conversion of fragment-level, raw count matrices to a bias- and expected background-corrected contact matrices of interaction scores. Pre-processing steps were performed prior to the post-processing steps of matrix balancing, binning, mean-variance relationship modeling and 5C replicate simulation. Binning and all subsequent normalization and modeling steps were performed on both experimental and simulated 5C replicates.

##### Data structure and pre-processing

We assembled sequencing counts from each 5C experiment $t_s$ and each genomic region $r$ into an $n_r \; x \; n_r$ raw contact matrix, $C_{t_s,r}$, where $n_r$ represents the total number of HindIII restriction fragments in each $r$, $t \in \{ES2i, ESserum, NPC\}$ represents a cellular condition, and $s \in \{1,2\}$ represents a biological replicate of the cellular condition $t$. Thus, $C_{t_s,r,i,j}$ is the number of reads that map to the $i^{th}$ and $j^{th}$ fragments in the $C_{t_s,r}$ matrix where $i \in \{1,2,3 \ldots n_r\}$ and $j \in \{1,2,3 \ldots n_r\}$. Raw contact matrices were then normalized as described[1]. Briefly, each $C_{t_s,r}$ was normalized for replicate biases due to batch effects, sequencing depth differences, and library complexity differences by conditional quantile normalization to create $C'_{t_s,r}$.

#### Matrix Balancing

Each $C'_{t_s,r}$ was then matrix balanced with joint express as described[1] to correct for differences in fragment-specific biases, such as GC content, fragment length, and 5C primer-specific efficiency at each primer in $r$ to create $C''_{t_s,r}$.

#### Contact matrix binning

Normalized and balanced contact matrices $C''_{t_s,r}$ were converted to interaction frequency matrices by binning at regular 4 kb intervals and smoothing at 16 kb intervals as described in[1, 3]. The smoothing was performed because we developed the 3DeFDR method on older 5C data from an alternating 5C primer design and the 3C template made in situ in the nucleus. The resulting binned interaction frequency matrices, $B_{t_s,r}$ , have $m_r$ by $m_r$ elements where $m_r$ is the total number of bins in the region. Thus, $B_{t_s,r,k,l}$ is computed as the arithmetic mean contact frequency between fragments in the $k^{th}$ and $l^{th}$ bins in genomic region $r$ as recorded in biological replicate $s$ under condition $t$. Binned interaction frequency values have reduced spatial noise relative to the original fragment-level counts while preserving the underlying structural signal.

#### Distance-dependence normalization

Following binning, expected values for each interaction in the binned interaction frequency matrices were computed using a modification of the local donut expected described by Aiden and colleagues that accounts for the local TAD/subTAD structure and the global distance-dependence background signal [1, 4]. The binned interaction frequency values (Observed), $B_{t_s,r,k,l}$ , were corrected by the maximum of expected donut values, $DE_{t_s,r,k,l}$ , and expected lower left values, $LLE_{t_s,r,k,l}$, to yield contact enrichments (Observed/Expected; Obs/Exp) normalized for distance-dependent 5C count signal and local chromatin domain structure.

#### Probabilistic model fitting

As detailed previously [1], contact enrichment values (Obs/Exp) were modeled within each region by parameterizing a log-logistic distribution using maximum likelihood estimation, resulting in p-value matrices denoted $P_{t_s,r}$. Right-tailed p-values were computed for each 5C genomic region separately.

**Removal of interactions below distance limit**

Interactions occurring between bins within 20 kb of each other on the linear chromatin fiber were removed from consideration and not included in further processing.

**Interaction scores**

The final step of the post-processing pipeline is the conversion of modeled p-values to interaction scores, denoted $IS_{t_s,r}$. For 3DeFDR, p-values were transformed to an interaction score of $-10 * log_2(pvalue)$. For benchmarking approaches ANOVA and 3DLRT (detailed below), p-values were transformed to interaction scores of $-10 * log_2(pvalue)$, as well as a z-score computed using the standard normal quantile function which is the inverse of the standard normal cumulative distribution function (**Equations 1 and 2**):

$$\Phi(P_{t_s,r,k,l}) = \frac{1}{\sqrt{2\pi}} \int_{-\infty}^{P_{t_s,r,k,l}} e^{-x^2/2} dx \quad \textbf{(1)}$$

$$Z_{t_s,r,k,l} = \Phi^{-1}(1 - P_{t_s,r,k,l}) \quad \textbf{(2)}$$

where $P_{t_s,r,k,l}$ is the right-tail p-value computed for the interaction between bins $k$ and $l$ in genomic region $r$ as recorded in biological replicate $s$ under condition $t$. We implemented the conversion of p-values to z-scores using the stats.norm.isf function in the scipy Python library.

**<u>3DeFDR</u>**

**Overview**

3DeFDR is designed to identify differential looping interactions across a set replicates containing at least two replicates in either two or three cellular conditions. In this section, we describe the application of 3DeFDR to three cellular conditions, referring to a set of three conditions $T = \{A, B, C\}$ and of six replicates as $S = \{A1, A2, B1, B2, C1, C2\}$. Our coding package is compatible with datasets of two or three conditions.

#### Differential loop categories

In the 3DeFDR framework, the set of possible classes of differential looping interactions is defined as all nonempty proper subsets, $H$, of the input condition set $T$ is (**Equation 3**):

$$H = \{\{A\}, \{B\}, \{C\}, \{A, B\}, \{A, C\}, \{B, C\}\} \quad \textbf{(3)}$$

Interactions assigned to single condition classes, e.g., $\{A\}$, $\{B\}$, or $\{C\}$ are considered to be interacting significantly higher in replicates of that specific condition than in those of the other two conditions (**Supplementary Figure 1**). Interactions assigned to dual condition classes, e.g. $\{A, B\}$, $\{A, C\}$, or $\{B, C\}$ are considered to be interacting significantly higher in replicates of the two specific conditions than in the remaining single condition (e.g. $C$, $B$, and $A$, respectively). If interaction scores are sufficiently high in all conditions, that interaction is interpreted to be non-differential and labeled as a constitutive looping interaction. If interaction scores are sufficiently low in all replicates of all conditions, the interaction is not called a looping interaction; therefore, it is not tested for differential looping signal.

#### Computing empirical false discovery rate

3DeFDR controls an empirically estimated false discovery rate (eFDR) to classify loops as differentially interacting across cellular condition set $T$. By definition, $FDR = E\left[\frac{V}{R}\right]$ where $V$ is the number of false positives among tests declared significant and $R$ is the total number of tests declared significant. $R$ is computed as the total number of pixels called as looping interactions with significantly differential interaction strength among all possible subsets, $H$, of the input condition set $T$. By contrast, $V$ is not trivially computed and requires a model for estimating what proportion of looping interactions in all possible subsets of H are false positives.

We hypothesized that $V$ is approximately equal to the total number of interactions incorrectly labeled differential when applying 3DeFDR to the set of biological samples known to have no truly differential loops, that is a null biological sample set. We defined our null data set as a set of samples that are replicates of a single cellular condition but assigned a set of labels matching conditions $T$. The key assumption of this approach is that that the false positive rate (FPR) of calls on the null dataset ($FPR_{null}$) is approximately equivalent to that of the experimental dataset ($FPR_{exp}$), such that $FPR_{null} \approx FPR_{exp}$. We computed and controlled an empirical false discovery rate (eFDR):

$$eFDR = \frac{n_{null}}{n_{exp}} \approx \frac{V}{R} \quad \textbf{(4)}$$

where $n_{exp}$ is the total number of interactions classified as significantly differential in the original experimental dataset and $n_{null}$ is the total number of interactions classified as significantly differential in the null dataset, which approximates $FPR_{exp.}$

We computed and controlled a piecewise eFDR thresholding scheme for each looping interaction class in the set of possible differential classifications $H$. To classify dynamic loops, 3DeFDR applies a thresholding scheme based on the difference in Interaction Scores between conditions (**Figure 1A-D**). Using a sweep of IS difference thresholds, $d$, (see orange lines in **Figure 1D**), 3DeFDR computes looping classifications for every pixel in a 5C data set and computes a class-specific eFDR for each differential looping interaction class ($h \in H$) as **(Equation 5)**:

$$eFDR_{d,h} \approx \frac{n_{null}^{d,h}}{n_{exp}^{d,h}} \quad \textbf{(5)}$$

where $n_{exp}^{d,h}$ is the total number of interactions assigned to differential looping class $h$ from the experimental replicates $S$ and $n_{null}^{d,h}$ is the total number of interactions assigned to differential looping class $h$ in the null replicates $S_{null}$ at difference threshold $d$. 3DeFDR adapts the distance threshold for each differential looping class to maintain a user-specified empirical FDR threshold $\tau$ across all differential looping classes. For each looping interaction class, $h$, we determined the lowest distance threshold at which

$eFDR_{d,h} \leq \tau$. We noticed that the relationship between eFDR and IS difference threshold $d$ is not always monotonically decreasing (**Supplementary Figure 3**), therefore we account for local minima in the curve prior to applying the FDR threshold $\tau$ to determine loop calls at the highest difference threshold $d$ at which the eFDR estimate is a minima that just passes the eFDR threshold. This parallels the explicit monotonization of BH-FDR corrected p-values. Thus, each differential looping class $h$ will have a unique difference threshold to reach a study-specific eFDR threshold $\tau$. Constitutive looping pixels are identified as those that pass the looping threshold but do not fall into any of the dynamic looping classes in $H$ (**Equation 3**). Overall, 3DeFDR employs FDR estimation control to guide the placement of IS thresholds to call differential looping classes.

We did not have access to an experimental dataset with enough replicates of the same cellular condition to create a null replicate set directly from real 5C libraries. To avoid the high costs and labor required to run additional experiments, we modeled and created simulations of our existing experimental replicates to create additional simulated replicates. We constructed a null dataset from the simulated replicates from one condition ($T_{null} = \{A, A, A\}$) and six replicates ($S_{null} = \{A1, A2, A3, A4, A5, A6\}$).

**Modeling and simulation of preprocessed replicates**

We simulated 5C replicates of the same condition at the level of fragment-level counts after conditional quantile normalization. The rationale for this decision is that library complexity, batch effect and sequencing depth terms would not have to be explicitly included in our models. Thus, we simulated fragment-resolution counts that have been quantile normalized (i.e. each $C'_{t_s,r,i,j}$ interaction) by fitting a negative binomial distribution to the counts across replicates of the same condition.

To begin constructing our simulation-generating model, we computed the sample mean and sample variance of the preprocessed sample counts of a single interaction across replicates of the same condition as in **Equations 6 and 7**:

$$\mu_{t,r,i,j} = \frac{\sum_{s=1}^{n_t} C'_{t_s,r,i,j}}{n_t} \quad \textbf{(6)}$$

$$\sigma^2{}_{t,r,i,j} = \frac{\sum_{s=1}^{n_t}\left(C'_{t_s,r,i,j} - \mu_{t_s,r,i,j}\right)^2}{n_t - 1} \tag{7}$$

where $n_t$is the number of replicates of condition $t,$ and $C'_{t_s,r,i,j}$ are the conditional quantile normalized raw 5C counts of the interaction between the $i^{th}$ and $j^{th}$bins of region $r$ in the $s^{th}$ replicate of condition $t$.

Most genomics experiments suffer from poor parameter estimation due to the low number of replicates that are financially and logistically feasible to generate for every biological condition. We did not use $\mu_{t,r,i,j}$ and $\sigma^2{}_{t,r,i,j}$ computed from $n_t = 2$ replicates to directly parameterize the negative binomial (NB) counts models for each $C'_{t_s,r,i,j}$ interaction. Rather, we modeled the mean-variance relationship (MVR) between $\mu_{t,r,i,j}$ and $\sigma^2{}_{t,r,i,j}$ by leveraging the high-dimensional nature of our data set to improve the $\sigma^2{}_{t,r,i,j}$ estimates. We stratified quantile normalized counts, $C'_{t_s,r,i,j}$ , for all regions by their linear genomic distance using an overlapping stratification windows of different sizes depending on genomic distance. For distance regime 1 (0-150 kb), we stratified the interactions using fine-grained, 12 kb-sized sliding windows, $w_{0\text{-}150kb}$, with a 4 kb step. For distance regime 2 (151-600 kb), we stratified the interactions into 24 kb-sized sliding windows, $w_{151\text{-}600kb}$, with an 8 kb step. For distance regime 3 (601-1000 kb), we stratified the interactions into coarse-grained, 60 kb-sized sliding windows, $w_{601\text{-}1000kb}$, with a 24 kb step. For each window $w$ in each distance regime, we modeled the MVR for each condition $t$ by fitting the function $\sigma^2 = A_{t,w}\mu^2 + \mu$ to the $\mu_{t,r,i,j}$ and $\sigma^2{}_{t,r,i,j}$ values for all regions $r$ and for all $i,j$ pairs whose linear genomic separation fell in window $w$. Prior to estimation of $A_{t,w}$, interactions with mean counts of one or less, or more than 2.5 standard deviations above the mean of mean counts for interactions in bin $w$ were removed. The dispersion parameters, $A_{t,w}$, were then plotted as a function of genomic distance, and Loess smoothing with a smoothing fraction of 0.5 was used to compute the final dispersion estimates, $\bar{A}_{t,w}$. The predicted sample variance for every window in all three distance regimes was then computed as in **Equation 8**:

$$\hat{\sigma}^2{}_{t,r,i,j} = \bar{A}_{t,w}\mu^2_{t,r,i,j} + \mu_{t,r,i,j} \tag{8}$$

where $w_{0\text{-}150kb}$ is a 12 kb-sized sliding window with a 4 kb step in distance regime 1, $w_{151\text{-}600kb}$ is a 24 kb-sized sliding window with a 8 kb step in distance regime 2, and $w_{601\text{-}100kb}$ is a 60 kb-sized sliding window with a 24 kb step in distance regime 3. We weighted the predicted variance value $\hat{\sigma}^2{}_{t,r,i,j}$ against the original observed variance of the interaction to generate a final weighted variance $\bar{\sigma}^2_{t,r,i,j}$ for each interaction in **Equation 9**:

$$\bar{\sigma}^2_{t,r,i,j} = \alpha\hat{\sigma}^2{}_{t,r,i,j} + \beta\sigma^2{}_{t,r,i,j} \qquad \textbf{(9)}$$

We chose to use $\alpha = \beta = 0.5$ to achieve pairwise correlations on par with that of real replicates while improving the quality of our variance estimate with the predicted contribution. As shown in **Supplementary Table 3**, increasing $\alpha$ led to higher pairwise correlation between simulated replicates. Finally, we parameterized the negative binomial model for each $C'_{t,r,i,j}$ interaction and generated simulated counts from it as in **Equation 10**:

$$C'^{sim}_{t,r,i,j} \sim NB(\mu_{t,r,i,j}, \bar{\sigma}^2_{t,r,i,j}) \qquad \textbf{(10)}$$

**Creating the null replicate set**

Using the generative models described above, we created six simulated replicates of each of the three cellular conditions, obtaining the simulated replicate set as in **Equation 11**:

$$S_{sim} = \{A1_{sim}, \ldots A6_{sim}, B1_{sim}, \ldots B6_{sim}, C1_{sim}, \ldots C6_{sim}\} \qquad \textbf{(11)}$$

These pseudoreplicates were then transformed in IS matrices using 5C processing pipeline described above. The user may then choose the replicate set of one condition (i.e. A, B, or C) for use as their null replicate set. For our results, we arbitrarily chose condition ES-2i.

**Identification of the background interaction set**

$C'^{sim}_{t_s,r,i,j}$ counts were then subject to the matrix balancing, binning, modeling, and p-value transformation steps described above. 3DeFDR takes as input the simulated replicate interaction scores, $IS^{sim}_{t_s,r}$, and experimental replicate interaction scores, $IS_{t_s,r}$, for each region $r$, replicate $s$, and condition $t$.

Prior to the identification of differential looping interactions, we created a background null interaction set if the interaction scores $IS_{t_s,r,k,l}$ of all replicates of every condition were less than a background threshold $b$ as in **Equation 12**:

$$Background\ loops\ =\ \{(r,k,l): \max_{t_s\ \in\ S}(IS_{t_s,r,k,l})\ <\ b\} \quad \textbf{(12)}$$

The exact threshold for background interactions that we used was $b = -10\times \log_2(0.8)$, corresponding to a p-value threshold of 0.8. Interactions not placed in this set were then passed on for further analysis for differential looping in the 3DeFDR pipeline.

#### Preliminary classification of differential looping interactions

As outlined in **Supplementary Figure 1**, to ultimately be classified as differential, a loop must pass thresholds for both baseline significance and IS difference across conditions.

#### Baseline significance filtering

To meet criteria for differential looping, for any differential classification $h$, an interaction must have $IS_{t_s,r,k,l}$ greater than a specific significance threshold $g$ for all replicates in at least one condition in $T$ as in **Supplementary Figure 1D** and **Equation 13**:

$$Significant\ Loops\ =\ \{(r,k,l): \max_t\left[\min_s\left(IS_{t_s,r,k,l}\right)\right] >\ g\} \quad \textbf{(13)}$$

In the current manuscript, the threshold for a significant looping interaction was $g = -10\times \log_2(0.165)$, corresponding to a p-value threshold of 0.165.

#### Thresholding interaction score differences across conditions

Starting with the subset of significant loops across conditions, we then set out to classify interactions according to how much their interaction scores changed across cellular conditions (**Supplementary Figure 1E)**. For each $(r, k, l)$ in the list of Significant Loops (**Equation 13**), we computed the difference in interaction score for every $IS_{t_s,r,k,l}$ interaction across each possible pair of replicates and conditions $t_s$. We then computed initial looping class assignments (**Equation 3)** across a sweep of IS difference thresholds $d$ as shown in **Supplementary Figure 1.** Additionally, in **Supplementary Table 2**, we provide the exact set of thresholds applied to obtain each possible looping classification of a bin-bin pair in dataset capturing three conditions.

In 3DeFDR, loop classifications are determined using this thresholding approach for each difference threshold across a sweep of all possible difference thresholds in a given data set. These classifications are considered preliminary prior to the application of the eFDR control procedure described in the next section.

**Final loop classification via Adaptive eFDR control procedure**

After obtaining preliminary classifications of each interaction across a sweep of IS difference thresholds, we determine each $IS_{r,k,l}$ interaction's final classification via the application of a classification-specific eFDR control procedure. For each possible loop classification $h \in H,$ we compute its eFDR for the sweep of difference threshold values, acquiring a difference threshold to eFDR mapping for each class, $eFDR_{d,h}$, as in **Equation 5**. We next apply the eFDR threshold $\tau$ to this mapping, isolating the highest difference threshold $d$ at which $eFDR_{d,h}$ is a minima that crosses below $\tau$ (**Supplementary Figure 3**), and report loop calls of class $h$ at this $d$. We perform the eFDR controlling procedure for every differential looping class $h \in H$ and the combined set of loop calls for each class constitutes our final set of differential classified loops.

Additionally, eFDR estimates can be computed as an average over a user-specified number, $N_{null-sets}$, of null replicate sets as in **Equation 14**:

$$eFDR_{d,h} = \frac{\frac{1}{N_{null-sets}} \sum_{m=1}^{N_{null-sets}} card\left(\left\{(r,k,l) \in h_{null}^{d}\right\}\right)}{card\left(\left\{(r,k,l) \in h_{exp}^{d}\right\}\right)} \qquad \textbf{(14)}$$

The numerator is now the average number of loops called as class $h$ in the null data sets at difference threshold $d$. The approach in **Equation 14** can reduce variability in eFDR estimates due to random differences between simulation sets generated from the same counts model.

**Benchmarking 3DeFDR on 5C**

To benchmark the effectiveness of 3DeFDR for loop calling, we implemented three additional methods for classifying differential looping interactions: (1) we applied conventional ANOVA, (2) we formulated a new likelihood ratio test, 3DLRT, and (3) we applied the published non-specific, Hi-C differential interaction caller diffHic on our 5C data. Using these three methods, we assigned a differential looping (DL) p-value to every interaction in an experimental dataset. In both approaches, output p-values were then corrected for multiple testing using the Benjamini-Hochberg step-up procedure for controlling FDR. Differential looping classifications were ultimately assigned using ANOVA and 3DLRT as detailed below and described in **Supplementary Figure 2**.

**ANOVA**

As a basic benchmark to compare to our more sophisticated methods, we applied ANOVA directly to either the Interaction Scores, $IS_{t_s,r,k,l}$, or the z-scores, $Z_{t_s,r,k,l}$, of the experimental replicate set. To account for the large number of interactions tested, we corrected the resulting differential interaction p-values for multiple testing by applying the Benjamini-Hochberg (BH) step-up procedure. The resulting BH FDR adjusted p-values were then thresholded according to a user-defined FDR to determine which interactions are significantly differential.

It should be noted that ANOVA may not be particularly appropriate when applied to the experimental design described in this paper. The Interaction Scores $IS_{t_s,r,k,l}$ are not normally distributed, whereas ANOVA assumes that the data it is applied to are normally distributed. Unlike the Interaction Scores, the z-scores $z_{t_s,r,k,l}$ are normally distributed with unit variance under the null hypothesis of the statistical model we use to call loops. Despite this, ANOVA attempts to independently re-estimate variance parameters for every $IS_{r,k,l}$ or $z_{t_s,r,k,l}$ interaction tested without leveraging the loop calling statistical model and without sharing information across interactions.

**3DLRT**

As an alternative method for identifying statistically significant differential interactions, we developed a new likelihood ratio test (3DLRT). 3DLRT compares the likelihood of the data assuming that the loop is not differentially interacting (null model) to the likelihood of the data under the assumption that the loop is differential (alternative model). We created the alternative model with parameters that best match the underlying data, whereas we created the null model with parameters that best match the null hypothesis. If the likelihood of the alternative model is significantly higher than the likelihood of the null model, then we reject the null hypothesis that the loop does not differentially change among conditions.

We formulated 3DLRT using either the z-scores, $Z_{t_s,r,k,l}$, (3DLRT-Z) or the Interaction Scores, $IS_{t_s,r,k,l}$, (3DLRT-IS) of the experimental replicate set. The derivation of 3DLRT that uses IS as input (3DLRT-IS) is provided in additional supplementary discussion below. For 3DLRT-Z, the z-scores are assumed to follow a unit-variance normal probability density function with a single looping effect size or shift parameter $\mu$ (**Equation 15**):

$$f_Z(Z;\mu) = \frac{1}{\sqrt{2\pi}} e^{-\frac{(Z-\mu)^2}{2}} \quad \textbf{(15)}$$

The test statistic for the likelihood ratio test based on the z-scores is shown as **Equation 16**:

$$T_Z = 2\ln \frac{\max_{\hat{\mu}_A,\hat{\mu}_B,\hat{\mu}_C}\left[\prod_{t\in T,s\in S} f_Z(Z_{t_s};\hat{\mu}_t)\right]}{\max_{\hat{\mu}_0}\left[\prod_{t\in T,s\in S} f_Z(Z_{t_s};\hat{\mu}_0)\right]} \quad \textbf{(16)}$$

where $T_Z$ is approximately Chi-squared distributed with two degrees of freedom. The intuition behind our formulation of 3DLRT is included below in the "Detailed description of the 3DLRT-Z test" section.

Finally, we assess the significance of the test statistic by comparing it to the chi-square distribution with degrees of freedom equal to the difference in the number of free parameters in our two models. In our case, the alternative hypothesis model has three parameters and the null hypothesis model has one, so we have two degrees of freedom. We then adjusted Chi-square p-values for multiple testing by applying the Benjamini-Hochberg step-up procedure. The resulting BH FDR adjusted p-values can then be thresholded according to a user-defined FDR to determine which interactions are significantly differential. Significantly differential interactions are then assigned classes using the same logic as 3DeFDR (**Supplementary Figure 4, Supplementary Table 2**).

**Detailed description of the 3DLRT-Z test**

**Set-up and assumptions**

The test statistic for 3DLRT-Z is shown in **Equation 17**:

$$T = 2\log\frac{\prod_i f(z_i;\hat{\mu}_{1i})}{\prod_i f(z_i;\hat{\mu}_{0i})} \quad \textbf{(17)}$$

where $f(x;\mu)$ is the probability density function for the normal distribution, constrained to unit variance and parameterized with a single looping effect size or shift parameter $\mu$, as specified in **(Equations 15-16)**, $\hat{\mu}_{0i}$ is the appropriate shift parameter estimate for $z_i$ under the null hypothesis, and $\hat{\mu}_{1i}$ is the appropriate shift parameter estimate for $z_i$ under the alternate hypothesis.

The parameter estimates $\vec{\hat{\mu}}_0$ and $\vec{\hat{\mu}}_1$ are constrained by the specific choice of null and alternate hypotheses. These parameters are chosen to maximize the likelihood of the data under the null (**Equation 18**) and alternate hypothesis (**Equation 19**), respectively:

$$\vec{\hat{\mu}}_0 = \text{argmax}_{\vec{\mu}_0} \prod_i f(z_i;\mu_{0i}) \text{, subject to null hypothesis constraints} \quad \textbf{(18)}$$

$$\vec{\hat{\mu}}_1 = \text{argmax}_{\vec{\mu}_1} \prod_i f(z_i;\mu_{1i}) \text{, subject to alternate hypothesis constraints} \quad \textbf{(19)}$$

Applying this choice of $\vec{\hat{\mu}}_0$ and $\vec{\hat{\mu}}_1$ to the equation for the likelihood ratio test statistic $T$ above, we derive **Equation 20**:

$$T = 2 \log \frac{\max_{\vec{\mu}_1} \prod_i f(z_i; \mu_{1i})}{\max_{\vec{\mu}_0} \prod_i f(z_i; \mu_{0i})} \quad \textbf{(20)}$$

**Equation 20** can also be written as **Equation 21**:

$$T = 2 \log \frac{\max_{\vec{\mu}_1} L(\vec{z} \mid \vec{\mu}_1)}{\max_{\vec{\mu}_0} L(\vec{z} \mid \vec{\mu}_0)} \quad \textbf{(21)}$$

where $L(\vec{z} \mid \vec{\mu}_1)$ is the total likelihood of the data $\vec{z}$ given a mean parameter vector $\vec{\mu}_1$ under the alternate hypothesis (**Equation 22)**:

$$L(\vec{z} \mid \vec{\mu}_1) = \prod_i f(z_i; \mu_{1i}) \quad \textbf{(22)}$$

and $L(\vec{z} \mid \vec{\mu}_0)$ is the total likelihood of the data $\vec{z}$ given a mean parameter vector $\vec{\mu}_0$ under the null hypothesis (**Equation 23)**:

$$L(\vec{z} \mid \vec{\mu}_0) = \prod_i f(z_i; \mu_{0i}) \quad \textbf{(23)}$$

The key difference between the two likelihood functions is that the constraints on $\vec{\mu}_1$ and $\vec{\mu}_0$ may be different, as dictated by the alternate and null hypotheses, respectively.

The likelihood ratio test statistic, $T$, will be assumed to follow a chi-square distribution under the null hypothesis. The degrees of freedom in the chi-square distribution will depend on the constraints imposed by the null and alternate hypotheses. For example, in our 3 condition, 2 replicate per condition experimental design, the parameters include (**Equations 24 and 25**):

$$\vec{\hat{\mu}}_1 = [\hat{\mu}_A, \hat{\mu}_A, \hat{\mu}_B, \hat{\mu}_B, \hat{\mu}_C, \hat{\mu}_C] \quad \textbf{(24)}$$

$$\vec{\hat{\mu}}_0 = [\hat{\mu}_{ABC}, \hat{\mu}_{ABC}, \hat{\mu}_{ABC}, \hat{\mu}_{ABC}, \hat{\mu}_{ABC}, \hat{\mu}_{ABC}] \quad \textbf{(25)}$$

and the chi-square distribution will have two degrees of freedom, since the alternate hypothesis has three free parameters, the null hypothesis has one free parameter, and $3 - 1 = 2$. More specifically, our null hypothesis is **Equation 26**:

$$\mu_A = \mu_B, \mu_A = \mu_C, \mu_B = \mu_C \quad \textbf{(26)}$$

and our alternate hypothesis is in **Equation 27**:

$$\mu_A \neq \mu_B \text{ OR } \mu_A \neq \mu_C \text{ OR } \mu_B \neq \mu_C \quad \textbf{(27)}$$

The single shift parameter for the null hypothesis is **Equation 28**:

$$\hat{\mu}_0 = \text{argmax}_\mu \prod_i f(x_i; \mu) = \hat{\mu}_{ABC} \quad \textbf{(28)}$$

The shift parameters for the alternate hypothesis are shown in **Equation 29**:

$$\hat{\mu}_{1i} = \begin{cases} \text{argmax}_\mu \prod_{i \in A} f(x_i; \mu) = \hat{\mu}_A & , \forall\, i \in A \\ \text{argmax}_\mu \prod_{i \in B} f(x_i; \mu) = \hat{\mu}_B & , \forall\, i \in B \\ \text{argmax}_\mu \prod_{i \in C} f(x_i; \mu) = \hat{\mu}_C & , \forall\, i \in C \end{cases} \quad \textbf{(29)}$$

For all six replicates in our three-condition experiment, the 3DLRT test statistic is then **Equation 30**:

$$T_{\text{LRT}} = 2 \log \frac{f(z_{A1}; \hat{\mu}_A) \times f(z_{A2}; \hat{\mu}_A) \times f(z_{B1}; \hat{\mu}_B) \times f(z_{B2}; \hat{\mu}_B) \times f(z_{C1}; \hat{\mu}_C) \times f(z_{C2}; \hat{\mu}_C)}{f(z_{A1}; \hat{\mu}_{ABC}) \times f(z_2; \hat{\mu}_{ABC}) \times f(z_{B1}; \hat{\mu}_{ABC}) \times f(z_{B2}; \hat{\mu}_{ABC}) \times f(z_{C1}; \hat{\mu}_{ABC}) \times f(z_{C2}; \hat{\mu}_{ABC})} \quad \textbf{(30)}$$

which can be rewritten more compactly to obtain **(Equation 16)**.

#### 3DLRT-Z p-value assignment and interaction classification

We call a single p-value for each bin-bin pair being tested for differential interaction strength from the likelihood ratio test statistic, $T_{\text{LRT}}$, using a chi-square distribution with two degrees of freedom, obtaining $P_{\text{LRT}}$ for each bin-bin pair. If there are $N$ bin-bin pairs being tested for differential interaction strength, there are $N$ of these p-values. We perform Benjamini-Hochberg multiple testing correction across all $N$ of these p-values, obtaining adjusted p-values $Q_{\mathrm{LRT}}$. Interactions whose $Q_{\mathrm{LRT}}$ is above the target false discovery rate are called "constitutive." Interactions whose $Q_{\mathrm{LRT}}$ is below the target false discovery rate are assigned a differential interaction classification category by first ranking the $\hat{\mu}$ values across the three conditions so that $\hat{\mu}_{A'} > \hat{\mu}_{B'} > \hat{\mu}_{C'}$, and then deciding between the $A'$ only and $A'B'$ classification categories by checking to see which of the pairs $(A', B')$ or $(B', C')$ have $\hat{\mu}$ values closer together. For example, if the pairwise comparisons have a low $Q_{\mathrm{LRT}}$, are assigned differential, and also follow $\hat{\mu}_{A'} > \hat{\mu}_{B'} > \hat{\mu}_{C'}$, then the classification category is assigned to be (**Equation 31**):

$$\text{classification category} = \begin{cases} A'\text{only} & , |\, \hat{\mu}_{A'} - \hat{\mu}_{B'}| > |\, \hat{\mu}_{B'} - \hat{\mu}_{C'}| \\ A'B' & , |\, \hat{\mu}_{A'} - \hat{\mu}_{B'}| < |\, \hat{\mu}_{B'} - \hat{\mu}_{C'}| \end{cases} \quad \textbf{(31)}$$

The full series of logic statements for all looping classifications are shown in **Supplementary Figure 4**.

**<u>Applying 3DeFDR and diffHic to genomic loci stratified from Hi-C data</u>**

To demonstrate the effectiveness of 3DeFDR for loop calling on Hi-C data, we modified our 5C pipeline to take Mb-sized genomic loci stratified from Hi-C libraries as input. We conducted a comparative analysis of differential looping classifications from 3DeFDR and diffHic [5] on the same region stratified from mouse ES cell and NPC Hi-C data (**Supplementary Table 5**).

Raw reads were processed with the Hi-C read alignment procedure detailed previously [7]. In brief, paired-end reads were aligned to mm9 mouse genome using Bowtie2 [8] through the HiC-Pro software [9]. Custom scripts were written to assemble Hi-C contact maps for each replicate at 10kb matrix resolution. We sliced ~3 Mb-sized genomic regions surrounding the *Sox2*, *Klf4*, *Olig1-2*, *Nanog, Nestin,* and *Synaptotagmin* genes from the two Hi-C maps. All downstream processing was performed using the 5C pipeline described above. Two specific modifications were made to the 5C pipeline: (1) we instituted conventional quantile normalization on the stratified Hi-C maps instead of the conditional quantile normalization procedure we conventionally employ for 5C data and (2) we did not use any smoothing given the high-resolution of the Hi-C data from Bonev et al. and the high complexity of the matrices at 10 kb matrix resolution. Similar to the 5C data, we used 3DeFDR to create sets of simulated null replicates from the stratified ES cell and NPC Hi-C matrices after quantile normalization. For this study, we used the ES cell condition to simulate null replicates. We converted simulated and real Hi-C matrices to interaction score matrices using the same 3DeFDR procedures described above.

3DeFDR thresholds were modified for use on Hi-C data. The background interaction threshold was $b = -10 \times \log_2(0.8)$, corresponding to a p-value threshold of 0.8, and the threshold for a significant looping interaction was $g = -10 \times \log_2(0.12)$, corresponding to a p-value threshold of 0.12. Differential loop classifications were then obtained with 3DeFDR using the same adaptive FDR control procedure described for 5C counts. Loops were reported at an eFDR of 30%.

### Applying diffHic to Hi-C and 5C data

We applied diffHiC following the package's provided usage guidelines for Hi-C data and applied a similar approach to run diffHic on 5C data.

#### *DiffHic on Hi-C*

We input our Hi-C dataset as raw, binned (10kb wide bins) counts. We floored these counts prior to inputting into diffHic. Library sizes were input as the total number of counts in each replicate. Following the diffHic user guide, we first filtered rows with any NaN counts. Then we filtered by average abundance (diffHic user guide section 4.1), discarding any bin-bin pair with an average abundance across all sample replicates of less than 5. We then used the direct filter (diffHic user guide section 4.2) to directly remove low-abundance bin-bin pairs, discarding any with abundances less than 2-fold higher than the estimated non-specific ligation rate (estimated as the median abundance across bin-bin pairs). We chose 2-fold above this estimate as our threshold instead of 5-fold, as presented in the usage instructions' example, because we found this threshold discarded too many points for us to perform later steps in the diffHiC pipeline. Next as recommended (diffHic user guide section 4.3), we filtered bin-bin pairs as a function of interaction distance, discarding pairs with abundances less than the value expected from compaction (generated using the filterTrended method, no parameters to set) choosing not to increase this threshold by an additional fold change value. Finally, we attempted to filter bin-bin pairs using the peak calling approach (diffHic user guide section 4.4). We found that this filter culled far too many points to allow us to run subsequent steps of the diffHic pipeline, regardless of the value of the peak width parameter (flank.width) while keeping other parameters at their values as recommended in the user guide (the minimum threshold for peak enrichment min.enrich at 0.5, the minimum threshold for peak counts min.count at 5, and the near diagonal cut-off min.diag of 2L). Next, we applied non-linear normalization (diffHic user guide section 5.2) to remove trended biases between libraries, specifically using LOESS normalization, using type="loess" for the normOffsets method. We also separately filtered near diagonal points, using by.dist=1.5e6 with method filterDiag and again LOESS normalization using type="loess" with method normOffSets as presented in the user guide (second half of diffHic section 5.2.1).

We then modeled biological variability between replicates. First, we estimated negative binomial dispersion for each bin-bin pair (diffHic user guide section 6.1-6.2) using a single-factor design matrix design = model.matrix(~conditions) where conditions is the corresponding list of the biological condition label for the input sample set. We performed estimation of negative binomial dispersion using this design. It is worth noting that the user manual indicates that major deviations from the monotonic trends might indicate batch effects in the data – and we did observe such deviations. We then estimated QL dispersion using this design as recommended in the user guide, setting robust=TRUE for method glmQLFit (diffHic user guide section 6.3).

Finally, we tested for significantly differential interactions using the quasi-likelihood F-test via method glmQLFTest (diffHic user guide section 7.1; no parameters to set) and obtained p-values adjusted to correct for multiple testing using the Benjamini-Hochberg (BH) method (diffHic user guide section 7.2). We then saved output log fold change and adjusted p-values for each bin-bin pair to file, and then visualized these results using our own tools. We thresholded bin-bin pairs to an FDR threshold, assigned loop classifications according to the sign of the log fold change value, and finally plotted these classifications spatially. Specifically, in **Supplementary Figure 13**, we ran applied this scheme with FDR thresholds of 0.3 (or 30%), 0.25, 0.2, 0.15, 0.1, 0.05, 0.01, and 0.001.

***DiffHic on 5C***

We input our 5C dataset as quantile normalized, binned (4kb wide bins), smoothed and matrix balanced counts. We floored these counts prior to inputting into diffHic. We then performed filtering and normalization steps of the diffHiC pipeline exactly as described above for Hi-C data. However, since our input dataset consisted of three biological conditions, we modified the remaining steps following recommendations in the edgeR user's guide for involving multiple pairwise comparisons involving more than two input treatment groups or conditions (edgeR user's guide section 3.2.3). We used the design matrix design=model.matrix(~conditions+0) and changed the column names of the design matrix to be simply the names of the conditions (ES2i, ESSerum, and NPC). We then estimated NB dispersions and QL dispersions as described above. Next we used makeContrasts to make the contrast vector for three comparisons across the three conditions. We compared each condition to the mean of the remaining two. We performed the F-test for each comparison, resulting in an associated log fold change and p-value for that comparison for each bin-bin pair. To assign a single differential p-value and a single possible looping classification to each bin-bin pair, we assigned an individual pair's p-value as the minimum across the three comparisons and then assigned its looping class according to the sign of the log fold change of the selected comparison. We then BH adjusted the selected p-values, and stored adjusted p-values and preliminary looping classifications to file. Using our own visualization tools, we again spatially plotted bin-bin pairs with adjusted p-values less than the FDR threshold of 1% (**Supplementary Figure 14**).

### SUPPLEMENTARY FIGURES

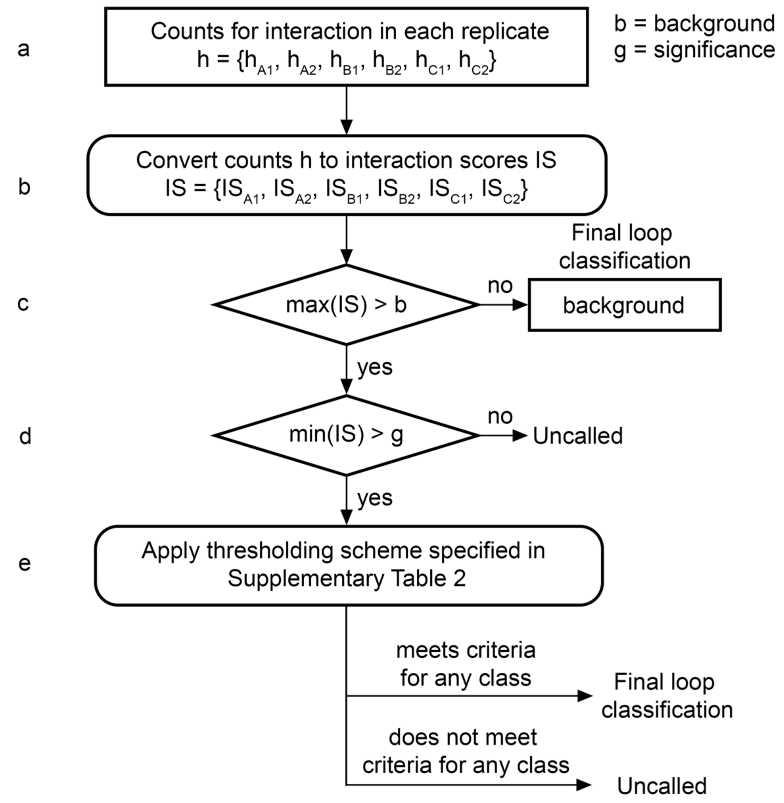


**Supplementary Figure 1. Overview schematic of 3DeFDR differential looping interaction classification procedure. (a)** For each conditionally quantile normalized fragment-fragment interaction, we have a set h of six sample counts values with two samples collected for each of three cellular conditions A, B, and C. **(b)** 5C counts are matrix balanced, binned, modeled, and converted to interaction scores (IS) (detailed in Supplementary Methods) and then subjected to a thresholding scheme. **(c)** If all sample IS values are lower than a certain background threshold value b, the bin-bin pair is labeled as a background loop. **(d)** To be tested for differential looping, a bin-bin pair must have at least one sample interaction score in set IS that is greater than a given significance threshold g. (A bin-bin pair that passes the background threshold control point but is not above the significance threshold in any condition will not be subject to further analysis and not assigned any label.) **(e)** We determine the final differential loop classification by thresholding differences in IS across conditions against the difference threshold and IS in each condition against the significance threshold, as specified in **Supplementary Table 2**. Final loop classifications are assigned based on the criteria listed **in Supplementary Table 2** with a loop being characteristic of a single condition (e.g. A only, B only, or C only) if it is significant in that condition and differential when compared to remaining conditions with respect to IS. Similarly, a loop is characteristic of two conditions (e.g. A & B, B & C, or A & C) if is significant in those conditions and differential when compared to the single remaining condition.

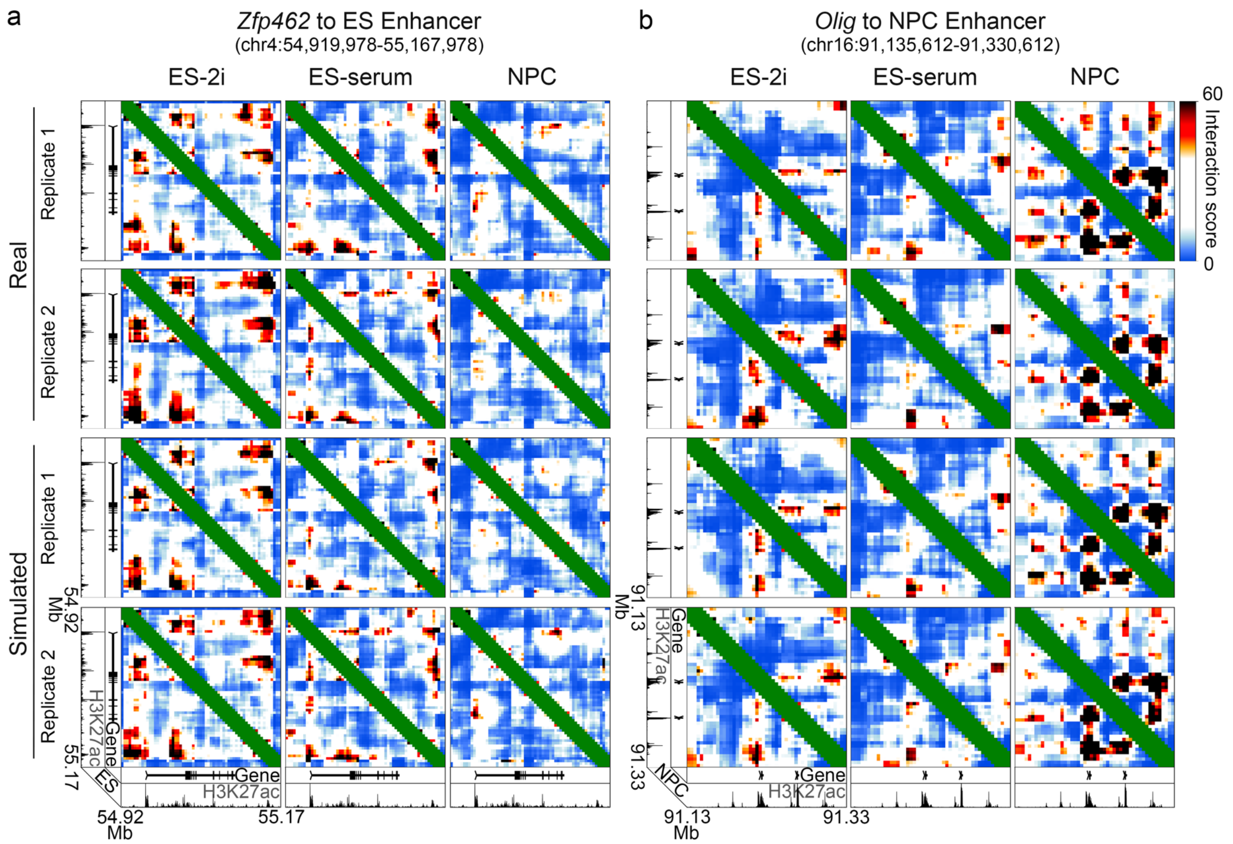


**Supplementary Figure 2**. **Real and simulated interaction score heatmaps for multiple replicates of each cell type. (a-b)** Interaction score heatmaps for real and simulated ES-2i, ES-serum, and NPC 5C replicates. (**a**) Zoomed-in heatmap encompassing *Zfp462* to ES-specific enhancer interaction. Putative enhancer demarcated by enriched ES-serum H3K27ac signal underneath the looping anchors. (**b**) Zoomed-in heatmap encompassing *Olig1/2* to NPC-specific enhancer interaction. Putative enhancer demarcated by enriched NPC H3K27ac signal underneath the looping anchors.

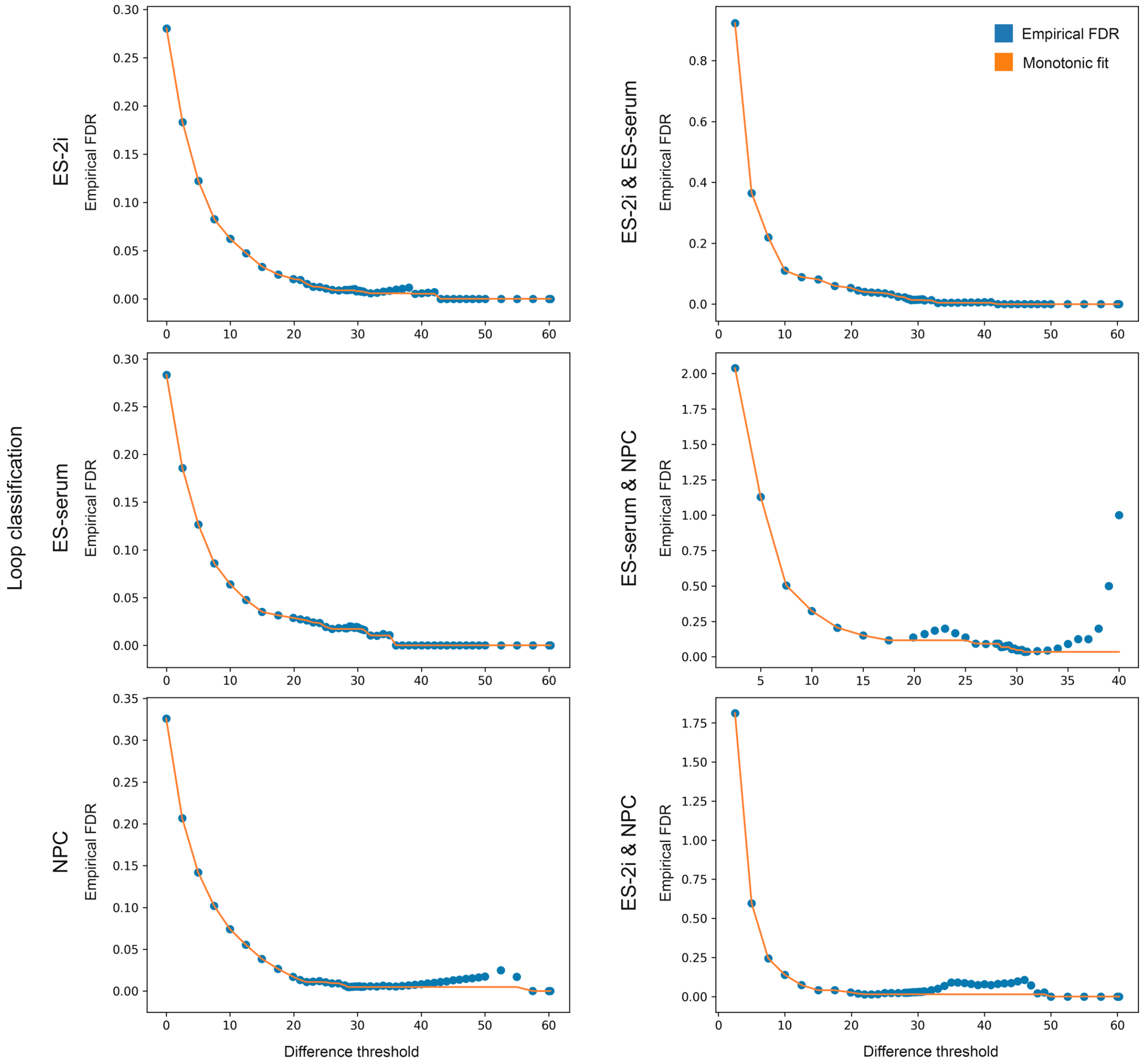


**Supplementary Figure 3. eFDR estimates across a sweep of difference thresholds for each possible differential loop classification when running 3DeFDR on the Beagan et al. 2017 5C dataset.** Each eFDR estimate vs. difference threshold trend (blue) is fitted with a monotonic minimum accumulator (orange) during 3DeFDR's eFDR control procedure to overcome noise in the estimate.

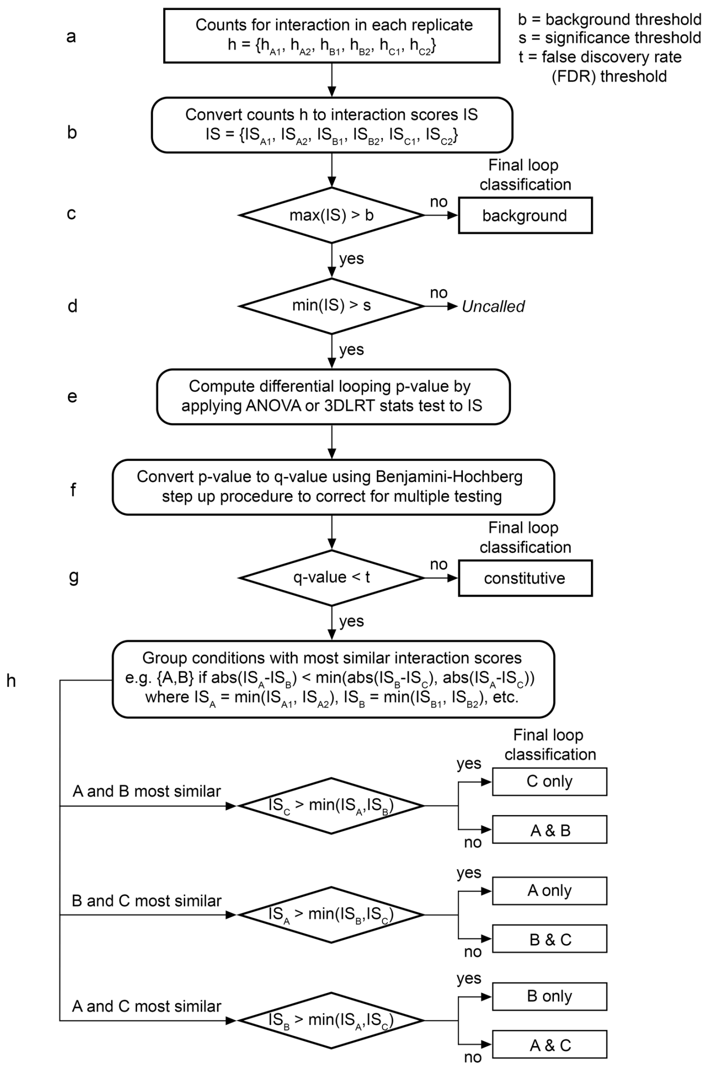


**Supplementary Figure 4. Overview schematic of ANOVA and 3DLRT differential looping interaction classification procedure. (a)** For each conditionally quantile normalized fragment-fragment interaction, we have a set h of six sample counts values with two samples collected for each of three cellular conditions A, B, and C. **(b)** 5C counts are matrix balanced, binned, modeled, and converted to interaction scores (IS), or alternatively z-scores, (detailed in Supplementary Methods) and then subjected to a thresholding scheme. **(c-d)** To be tested for differential looping, a bin-bin pair must have at least one sample interaction score in set IS that is greater than a given significance threshold. Additionally, if all sample interaction scores in IS are lower than a specific background threshold value, the bin-bin pair is labeled as a background loop. A bin-bin pair that passes the background threshold control point but is not above the significance threshold in any condition will not be subject to further analysis and not assigned any label. **(e-f)** To identify a bin-bin pair as significantly differential across cellular conditions, we first compute its differential looping p-value using either the ANOVA or 3DLRT statistical test. To account for multiple testing, p-values are adjusted to q-values with Benjamini-Hochberg. **(g)** If the resulting differential looping q-value is lower than a user-defined false discovery rate (FDR) threshold t, the interaction is classified as differential and otherwise is classified as non-differential or constitutive across conditions. **(h)** Differential interactions are further categorized according to the direction and fold change of their differential looping signal to obtain a final looping classification.

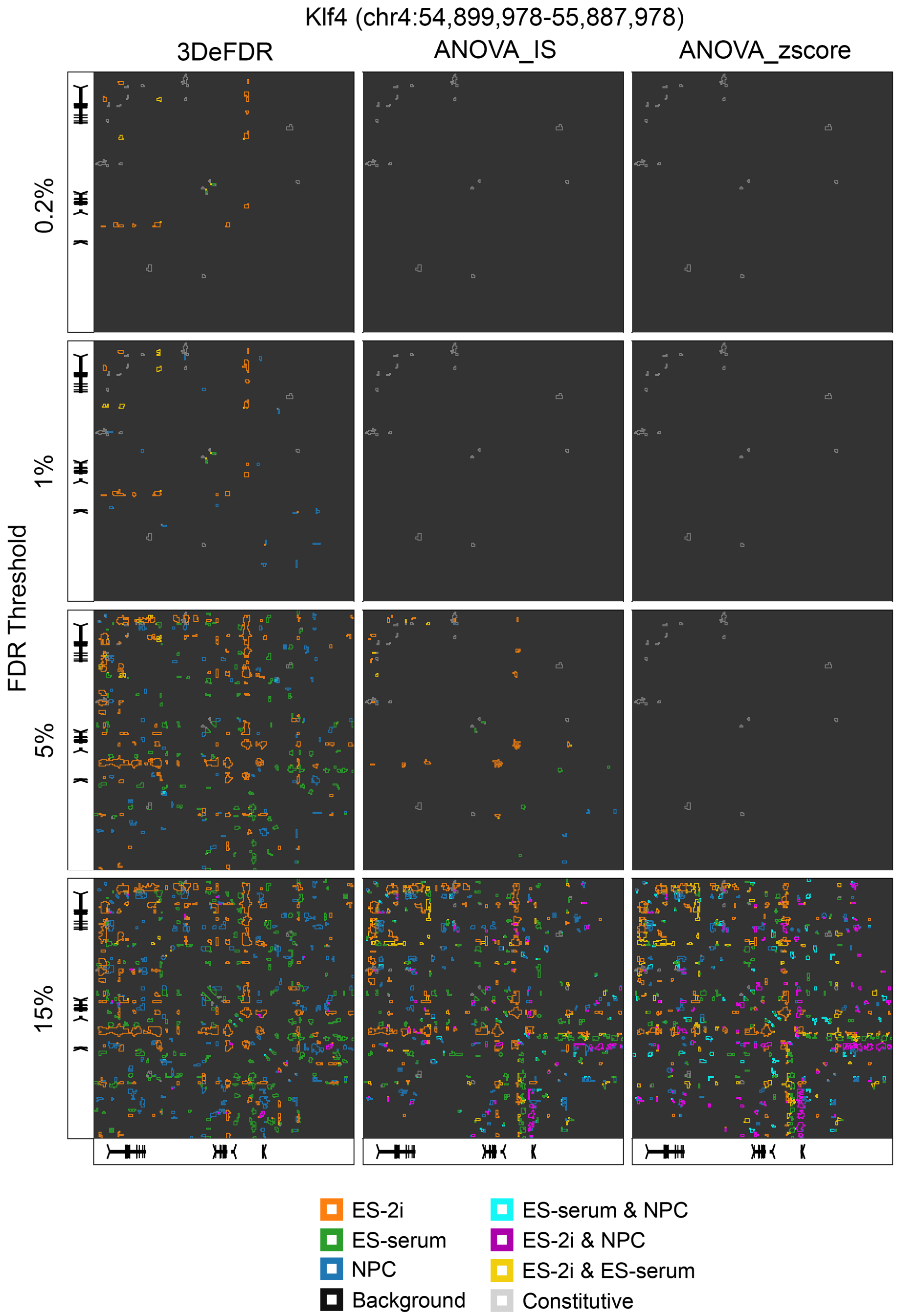


**Supplementary Figure 5. Benchmarking of 3DeFDR at the *Klf4* locus against ANOVA tests performed on interaction scores and z-scores.** Looping interaction classes identified in the genomic region surrounding *Klf4* via 3DeFDR, ANOVA on interaction scores, and ANOVA on z-scores across a sweep of false discovery rate (FDR) thresholds. All three approaches used the interaction score as the random variable (Supplementary Methods).

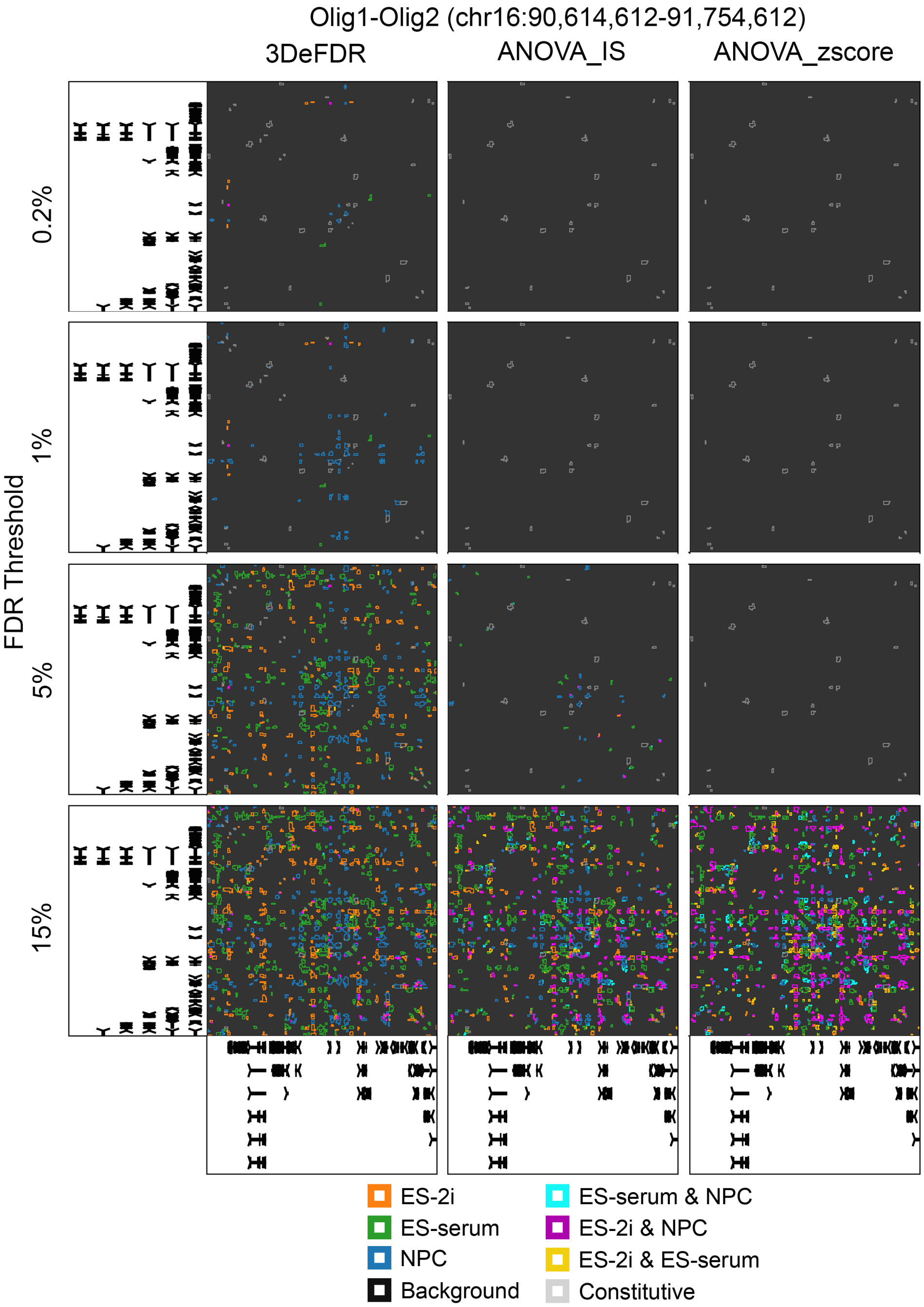


**Supplementary Figure 6. Benchmarking of 3DeFDR at the *Olig1-Olig2* locus against ANOVA tests performed on interaction scores and z-scores.** Looping interaction classes identified in the genomic region surrounding *Olig1/2* via 3DeFDR, ANOVA on interaction scores, and ANOVA on z-scores across a sweep of false discovery rate (FDR) thresholds. All three approaches used the interaction score as the random variable (Supplementary Methods).

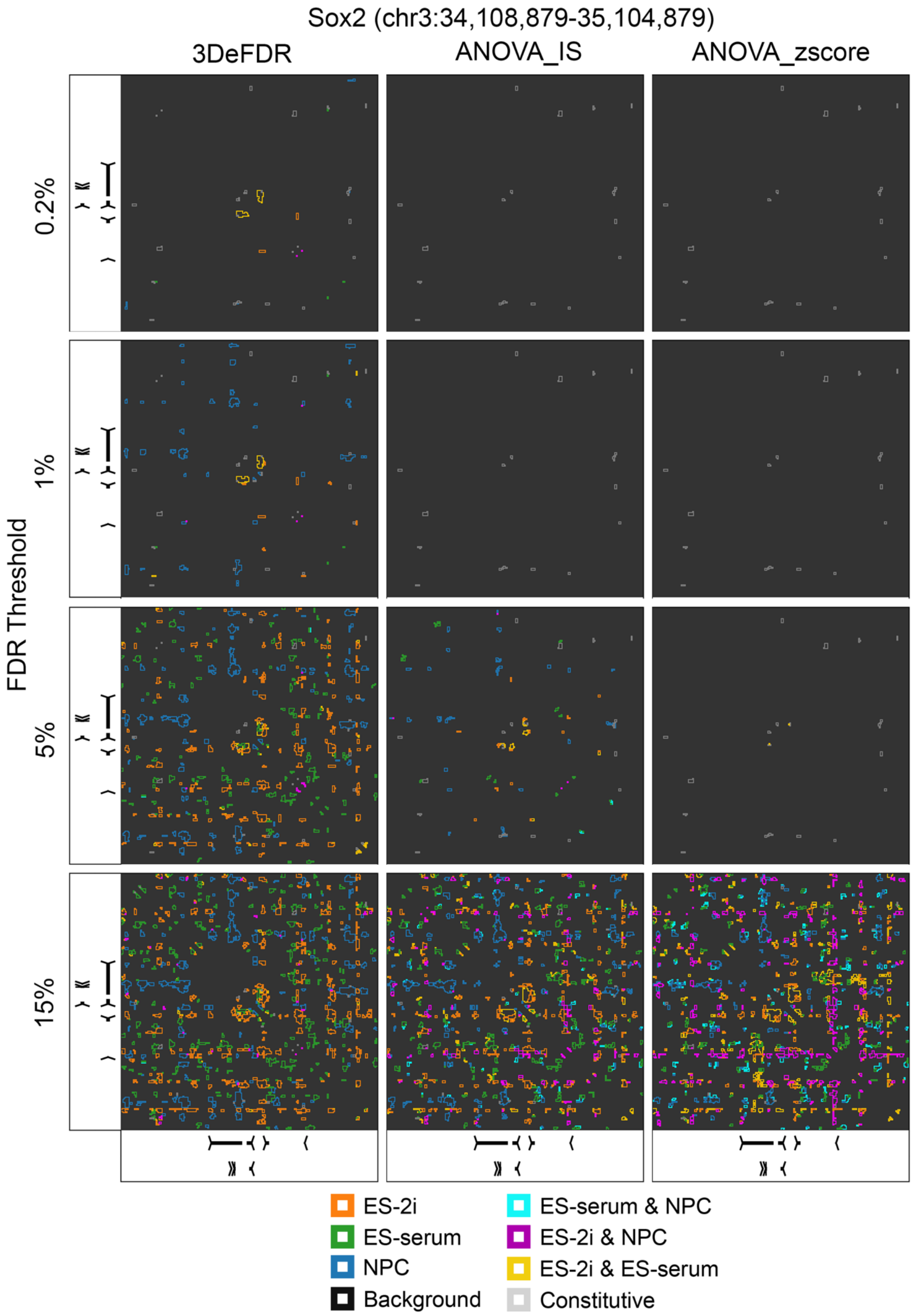


**Supplementary Figure 7. Benchmarking of 3DeFDR at the *Sox2* locus against ANOVA tests performed on interaction scores and z-scores.** Looping interaction classes identified in the genomic region surrounding *Sox2* via 3DeFDR, ANOVA on interaction scores, and ANOVA on z-scores across a sweep of false discovery rate (FDR) thresholds. All three approaches used the interaction score as the random variable (Supplementary Methods).

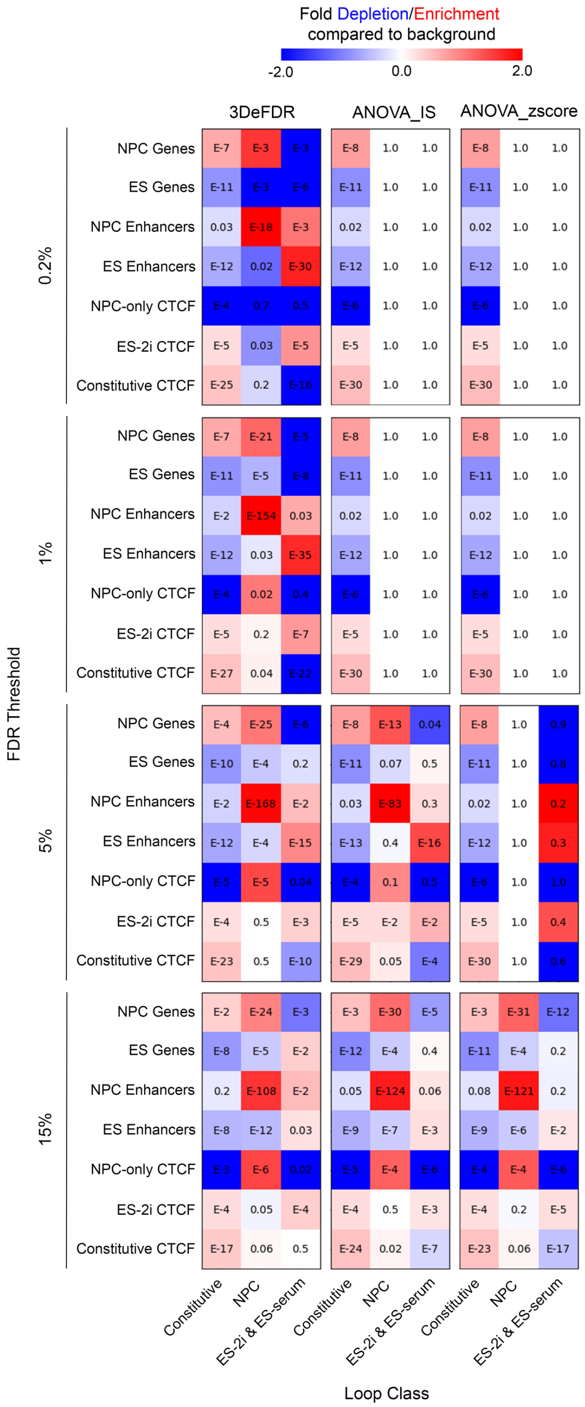


**Supplementary Figure 8. Fold change depletion/enrichment in cell-type specific chromatin features for differential loops classified via 3DeFDR, ANOVA performed on interaction scores, and ANOVA performed on z-scores.** Fold change depletion/enrichment is computed as the proportion of binned interactions assigned to a given looping class (columns) that have at least one anchor occupied by a given chromatin feature (rows) over the proportion of binned interactions labeled background which are likewise positive for that feature. P-values included within each entry are computed using Fischer's exact test.

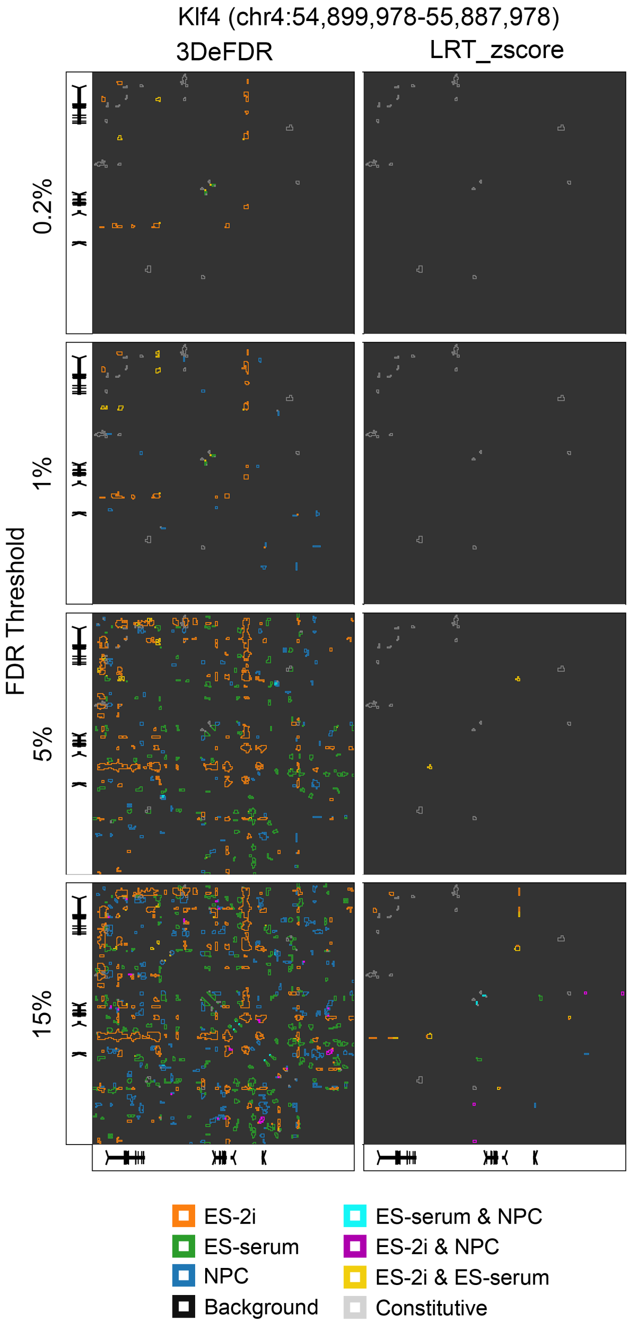


**Supplementary Figure 9. Benchmarking of 3DeFDR at the *Klf4* locus against 3DLRT performed on z-scores.** Looping interaction classes identified in the genomic region surrounding *Klf4* via 3DeFDR and 3DLRT on z-scores across a sweep of false discovery rate (FDR) thresholds. All three approaches used the interaction score as the random variable (Supplementary Methods).

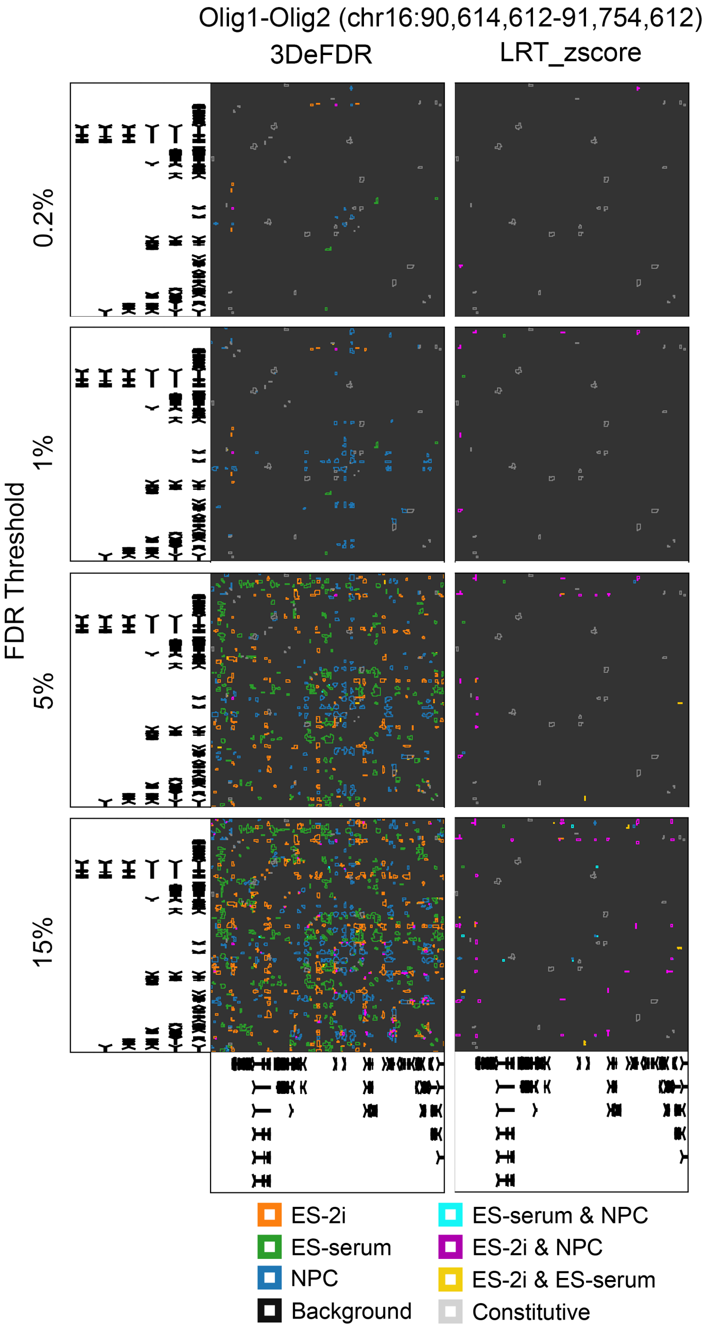


**Supplementary Figure 10. Benchmarking of 3DeFDR at the *Olig1-Olig2* locus against 3DLRT performed on z-scores.** Looping interaction classes identified in the genomic region surrounding *Olig1/2* via 3DeFDR and 3DLRT on z-scores across a sweep of false discovery rate (FDR) thresholds. All three approaches used the interaction score as the random variable (Supplementary Methods).

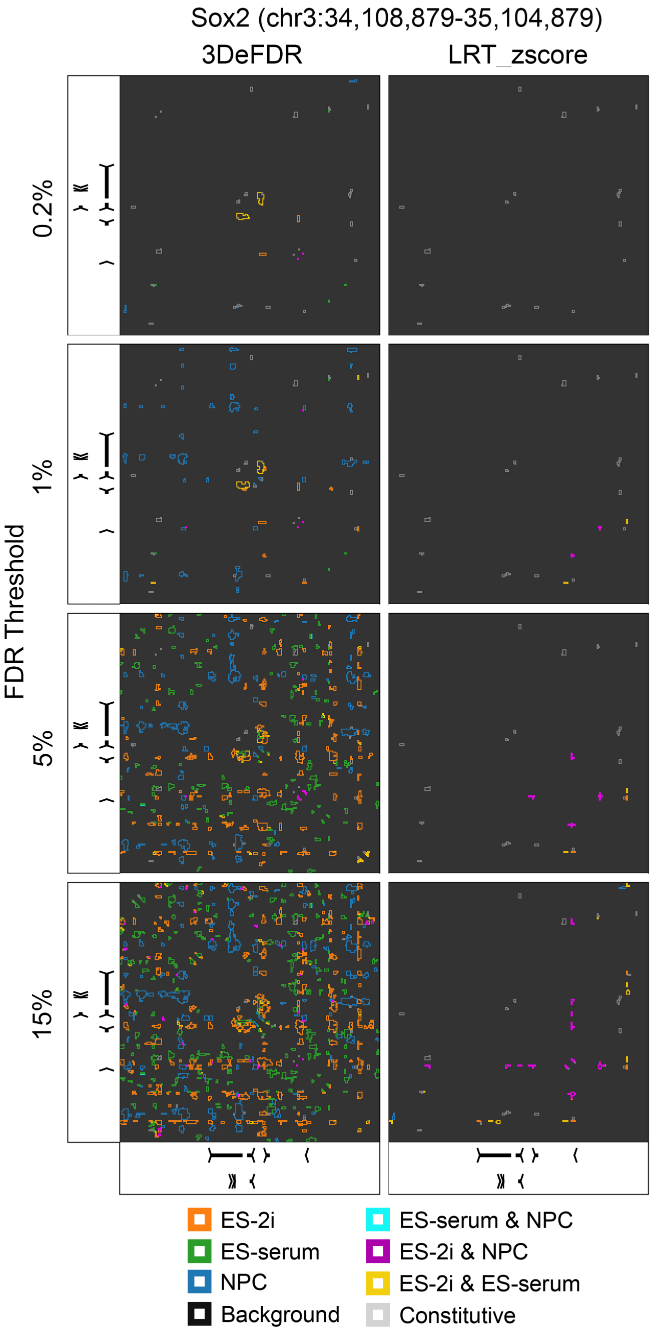


**Supplementary Figure 11. Benchmarking of 3DeFDR at the *Sox2* locus against 3DLRT performed on and z-scores.** Looping interaction classes identified in the genomic region surrounding *Sox2* via 3DeFDR and 3DLRT on z-scores across a sweep of false discovery rate (FDR) thresholds. All three approaches used the interaction score as the random variable (Supplementary Methods).

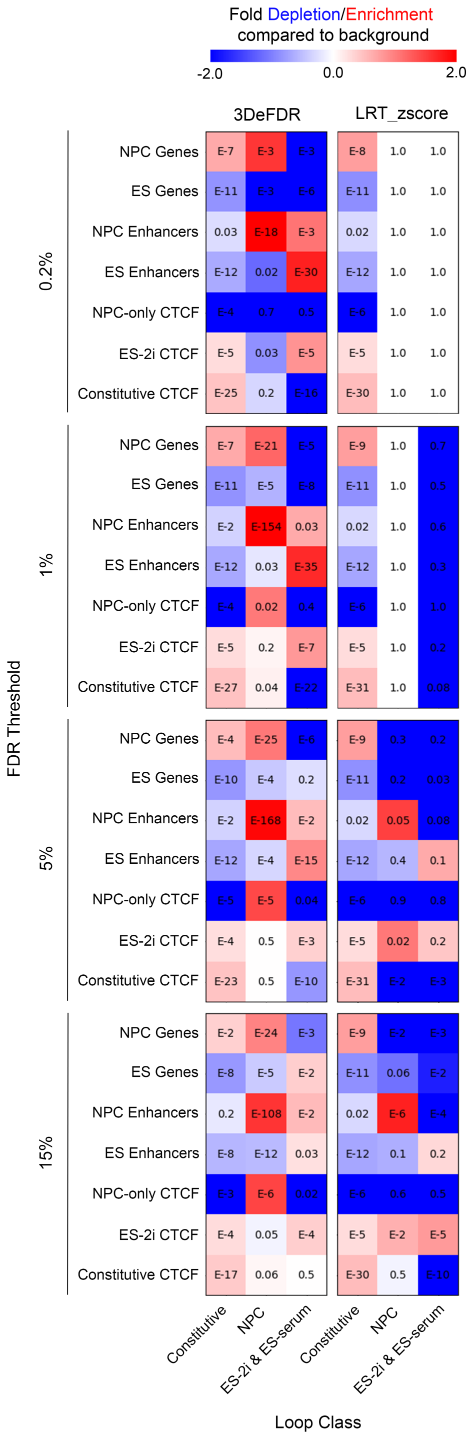


**Supplementary Figure 12. Fold change depletion/enrichment in cell-type specific chromatin features for differential loops classified via 3DeFDR and 3DLRT performed on z-scores.** Fold change depletion/enrichment is computed as the proportion of binned interactions assigned to a given looping class (columns) that have at least one anchor occupied by a given chromatin feature (rows) over the proportion of binned interactions labeled background which are likewise positive for that feature. P-values included within each entry are computed using Fischer's exact test.

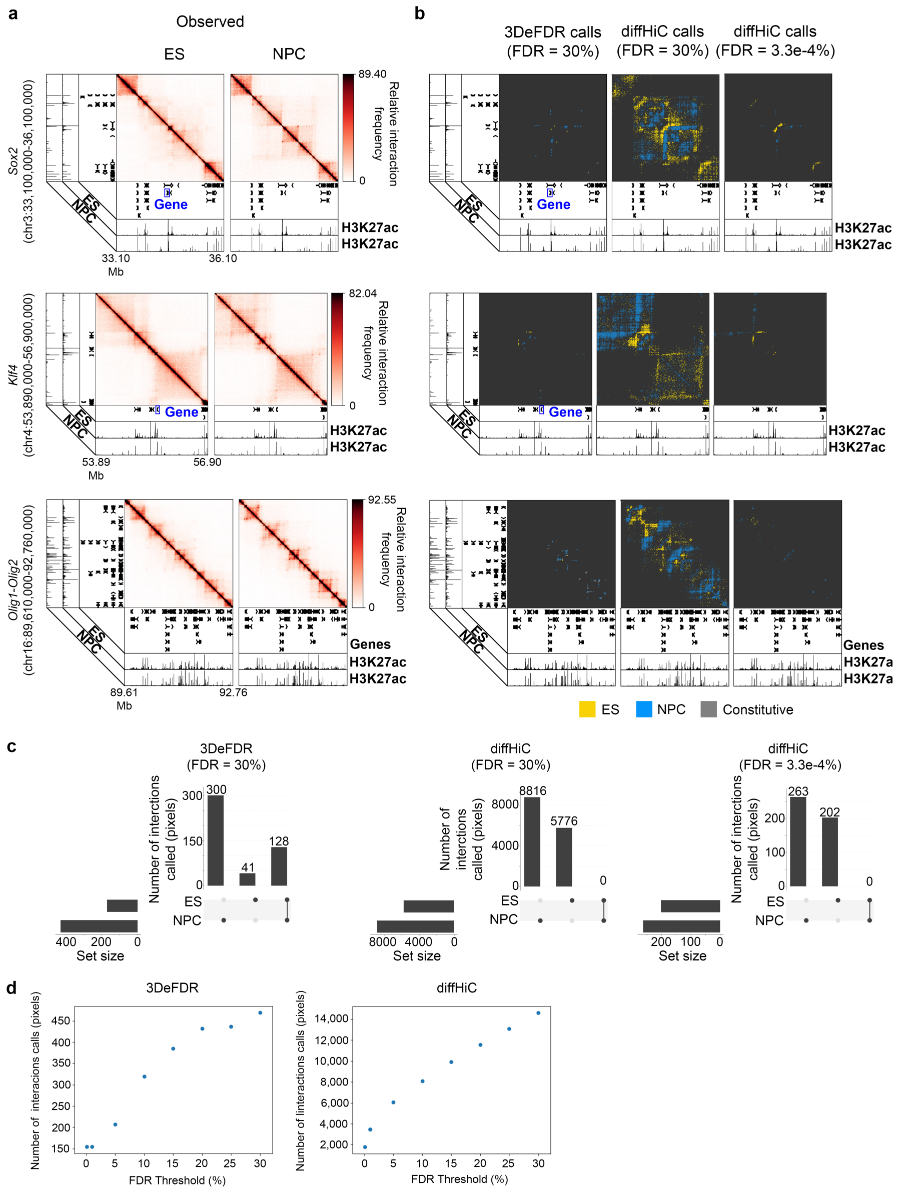


**Supplementary Figure 13. Differential loops classified from Hi-C data via 3DeFDR and diffHiC. (a)** Reference heatmaps representing binned, quantile normalized, and joint express normalized Hi-C counts (Observed) around the *Sox2* (chr3:33,100,000-36,100,000), *Klf4* (chr4:53,890,000-56,900,000), *Olig1-Olig2* (chr16:89,610,000-92,760,000) genes. **(b)** Looping interaction classes identified across 6 genomic regions cut from a Hi-C dataset. Loops are called using 3DeFDR with an FDR threshold of 30% (469 pixels called in the 3 regions shown), diffHiC at a matched FDR threshold of 30%, and diffHiC at an FDR threshold that results in the same total number of differentially classified pixels across the three regions as 3DeFDR at an FDR threshold of 30% (465 pixels at an FDR threshold of 3.3e-4%). **(c)** UpSetR scalable Venn diagrams of the number of pixels called as ES only, NPC only, or constitutive. **(d)** Scatterplot of number of pixels called as differential across a sweep of FDR thresholds for 3DeFDR and diffHiC across the three regions shown.

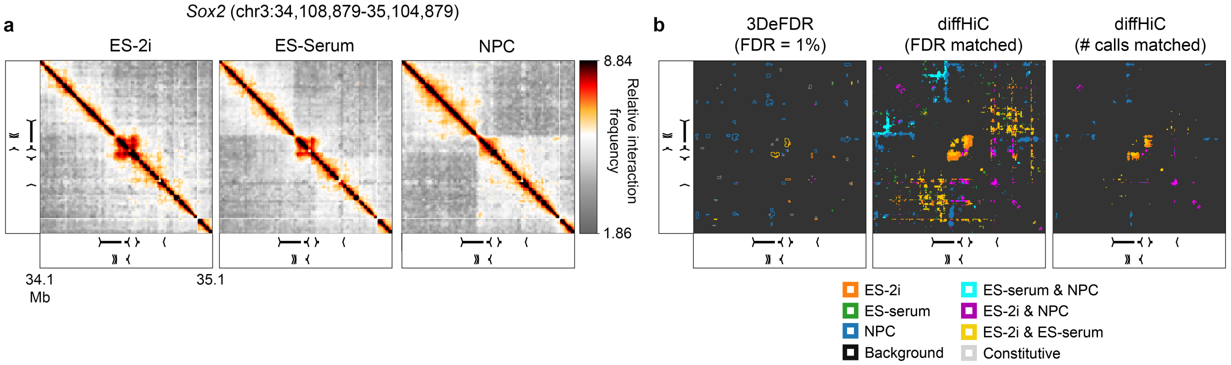


**Supplementary Figure 14. Differential loops classified from 5C data via 3DeFDR and diffHiC. (a)** Reference heatmaps representing quantile normalized, joint express normalized, and binned 5C counts (Observed) around the *Sox2* gene (chr3:34,108,879-35,104,879) **(b)** Looping interaction classes identified in each genomic region using 3DeFDR with FDR threshold = 1%, diffHiC using the same FDR threshold, and diffHiC using an FDR threshold that results in the same total number of differentially classified pixels as 3DeFDR at its reported FDR threshold. In this region alone, the total number of pixels called by 3DeFDR at FDR = 1% is 454 and for diffHiC at 0.07% is 456.

**SUPPLEMENTARY TABLES**

**Supplementary Table 1. Publicly available 5C libraries.**

| **Cell Type** | **Sample** | **Reference** | **Accession number** |
|---|---|---|---|
| mES (V6.5) in 2i/LIF media | ES 2i, Rep 1 | Beagan et al. 2017 | GSM2259911 |
| mES (V6.5) in 2i/LIF media | ES 2i, Rep 2 | Beagan et al. 2017 | GSM2259912 |
| mES (V6.5) in serum/LIF media | ES Serum, Rep 1 | Beagan et al. 2017 | GSM2259913 |
| mES (V6.5) in serum/LIF media | ES Serum, Rep 2 | Beagan et al. 2017 | GSM2259914 |
| Primary NPCs | NPC, Rep 1 | Beagan et al. 2017 | GSM2259915 |
| Primary NPCs | NPC, Rep 2 | Beagan et al. 2017 | GSM2259916 |

**Supplementary Table 2. 3DeFDR criteria for differential loop classification.**

| **Looping class** | **Significance criteria** ($g$ significance threshold) | **Differential looping criteria** ($d$ interaction score difference threshold) |
|---|---|---|
| {A} | $IS_{A1} > g$<br>$IS_{A2} > g$ | $IS_{A1} - IS_{B1} > d$<br>$IS_{A1} - IS_{B2} > d$<br>$IS_{A2} - IS_{B1} > d$<br>$IS_{A2} - IS_{B2} > d$<br><br>$IS_{A1} - IS_{C2} > d$<br>$IS_{A2} - IS_{C1} > d$<br>$IS_{A2} - IS_{C2} > d$ |
| {B} | $IS_{B1} > g$<br>$IS_{B2} > g$ | $IS_{B1} - IS_{A1} > d$<br>$IS_{B1} - IS_{A2} > d$<br>$IS_{B2} - IS_{A1} > d$<br>$IS_{B2} - IS_{A2} > d$<br><br>$IS_{B1} - IS_{C1} > d$<br>$IS_{B1} - IS_{C2} > d$<br>$IS_{B2} - IS_{C1} > d$<br>$IS_{B2} - IS_{C2} > d$ |
| {C} | $IS_{C1} > g$<br>$IS_{C2} > g$ | $IS_{C1} - IS_{A1} > d$<br>$IS_{C1} - IS_{A2} > d$<br>$IS_{C2} - IS_{A1} > d$<br>$IS_{C2} - IS_{A2} > d$<br><br>$IS_{C1} - IS_{B1} > d$<br>$IS_{C1} - IS_{B2} > d$<br>$IS_{C2} - IS_{B1} > d$<br>$IS_{C2} - IS_{B2} > d$ |

| | | |
|---|---|---|
| {A, B} | $IS_{A1} > g$<br>$IS_{A2} > g$<br><br>$IS_{B1} > g$<br>$IS_{B2} > g$ | $IS_{A1} - IS_{C1} > d$<br>$IS_{A1} - IS_{C2} > d$<br>$IS_{A2} - IS_{C1} > d$<br>$IS_{A2} - IS_{C2} > d$<br><br>$IS_{B1} - IS_{C1} > d$<br>$IS_{B1} - IS_{C2} > d$<br>$IS_{B2} - IS_{C1} > d$<br>$IS_{B2} - IS_{C2} > d$<br><br>$\|IS_{A1} - IS_{B1}\| \leq d$<br>$\|IS_{A1} - IS_{B2}\| \leq d$<br>$\|IS_{A2} - IS_{B1}\| \leq d$<br>$\|IS_{A2} - IS_{B2}\| \leq d$ |
| {A, C} | $IS_{A1} > g$<br>$IS_{A2} > g$<br><br>$IS_{C1} > g$<br>$IS_{C2} > g$ | $IS_{A1} - IS_{B1} > d$<br>$IS_{A1} - IS_{B2} > d$<br>$IS_{A2} - IS_{B1} > d$<br>$IS_{A2} - IS_{B2} > d$<br><br>$IS_{C1} - IS_{B1} > d$<br>$IS_{C1} - IS_{B2} > d$<br>$IS_{C2} - IS_{B1} > d$<br>$IS_{C2} - IS_{B2} > d$<br><br>$\|IS_{A1} - IS_{C1}\| \leq d$<br>$\|IS_{A1} - IS_{C2}\| \leq d$<br>$\|IS_{A2} - IS_{C1}\| \leq d$<br>$\|IS_{A2} - IS_{C2}\| \leq d$ |
| {B, C} | $IS_{B1} > g$<br>$IS_{B2} > g$<br><br>$IS_{C1} > g$<br>$IS_{C2} > g$ | $IS_{B1} - IS_{A1} > d$<br>$IS_{B1} - IS_{A2} > d$<br>$IS_{B2} - IS_{A1} > d$<br>$IS_{B2} - IS_{A2} > d$<br><br>$IS_{C1} - IS_{A1} > d$<br>$IS_{C1} - IS_{A2} > d$<br>$IS_{C2} - IS_{A1} > d$ |

| | | |
|---|---|---|
| | | $IS_{C2} - IS_{A2} > d$<br><br>$\|IS_{B1} - IS_{C1}\| \leq d$<br>$\|IS_{B1} - IS_{C2}\| \leq d$<br>$\|IS_{B2} - IS_{C1}\| \leq d$<br>$\|IS_{B2} - IS_{C2}\| \leq d$ |
| Constitutive | $IS_{A1} > g$<br>$IS_{A2} > g$<br><br>$IS_{B1} > g$<br>$IS_{B2} > g$<br><br>$IS_{C1} > g$<br>$IS_{C2} > g$ | $\|IS_{A1} - IS_{B1}\| \leq d$<br>$\|IS_{A1} - IS_{B2}\| \leq d$<br>$\|IS_{A2} - IS_{B1}\| \leq d$<br>$\|IS_{A2} - IS_{B2}\| \leq d$<br><br>$\|IS_{A1} - IS_{C1}\| \leq d$<br>$\|IS_{A1} - IS_{C2}\| \leq d$<br>$\|IS_{A2} - IS_{C1}\| \leq d$<br>$\|IS_{A2} - IS_{C2}\| \leq d$<br><br>$\|IS_{B1} - IS_{C1}\| \leq d$<br>$\|IS_{B1} - IS_{C2}\| \leq d$<br>$\|IS_{B2} - IS_{C1}\| \leq d$<br>$\|IS_{B2} - IS_{C2}\| \leq d$ |

* We use the shorthand $IS_{A1}$ to refer to the sample interaction score of a single loop between bins *k* and *l* in region *r* for replicate 1 of condition A, a value denoted $IS_{A_1,r,k,l}$ in the Supplementary Methods.

**Supplementary Table 3. Spearmann's correlations between simulated and real 5C counts data.**

| Mean correlation between real replicates | | |
|---|---|---|
| ES-2i | ES-Serum | NPC |
| 0.925 | 0.933 | 0.885 |

| Interreplicate variance weight | | Mean correlation between simulated replicates | | |
|---|---|---|---|---|
| **Predicted** | **Observed** | **ES-2i** | **ES-Serum** | **NPC** |
| 0.00 | 1.00 | 0.929 | 0.933 | 0.903 |
| 0.25 | 0.75 | 0.940 | 0.942 | 0.917 |
| 0.50 | 0.50 | 0.947 | 0.949 | 0.926 |
| 0.75 | 0.25 | 0.953 | 0.953 | 0.934 |
| 1.00 | 0.00 | 0.959 | 0.958 | 0.943 |
| 1.00 | 1.00 | 0.912 | 0.916 | 0.880 |
| 0.00 | 2.00 | 0.892 | 0.900 | 0.857 |
| 2.00 | 0.00 | 0.923 | 0.921 | 0.900 |

### Supplementary Table 4. Publicly available ChIP-seq datasets.

| | | | | | |
|---|---|---|---|---|---|
| **Target protein** | CTCF | CTCF | CTCF | H3K27ac | H3K27ac |
| **Cell type** | Mouse ES cells (V6.5) in 2i/LIF media | Mouse ES cells (V6.5) in serum/LIF media | Primary NPCs | Mouse ES cells (V6.5) | ES-derived NPC |
| **Mapped test ChIP-seq reads after downsampling** | 11000000 | 11000000 | 11000000 | 7000000 | 7000000 |
| **Test ChIP-seq reference** | Beagan et al. 2017 | Beagan et al. 2017 | Beagan et al. 2017 | Creyghton et al. 2010 | Creyghton et al. 2010 |
| **Test sample GEO accession number** | GSM2259905 | GSM2259907 | GSM2259909 | GSM594579 | GSM594585 |
| **Control samples** | Mouse ES cells (V6.5) in 2i/LIF whole cell extract | Mouse ES cells (V6.5) in serum/LIF whole cell extract | Primary NPCs whole cell extract | V6.5 whole cell extract | NPC whole cell extract |
| **Mapped control ChIP-seq reads after downsampling** | 15000000 | 15000000 | 15000000 | 7000000 | 7000000 |
| **Control sample GEO accession number** | GSM2259906 | GSM2259908 | GSM2259910 | GSM594599 | GSM883648 |

**Supplementary Table 5. Publicly available Hi-C libraries.**

| Cell Type | Sample | Reference | Accession number |
|---|---|---|---|
| Mouse ES cells | ES Serum, Rep 1 | Bonev et al. 2017 | GSM2533818 |
| Mouse ES cells | ES Serum, Rep 2 | Bonev et al. 2017 | GSM2533819 |
| Mouse ES cells | ES Serum, Rep 3 | Bonev et al. 2017 | GSM2533820 |
| Mouse ES cells | ES Serum, Rep 4 | Bonev et al. 2017 | GSM2533821 |
| In vitro differentiated NPCs | NPC, Rep 1 | Bonev et al. 2017 | GSM2533822 |
| In vitro differentiated NPCs | NPC, Rep 2 | Bonev et al. 2017 | GSM2533823 |
| In vitro differentiated NPCs | NPC, Rep 3 | Bonev et al. 2017 | GSM2533824 |
| In vitro differentiated NPCs | NPC, Rep 4 | Bonev et al. 2017 | GSM2533825 |
